# Pretransplant targeting of TNFRSF25 and CD25 stimulates recipient Tregs in target tissues ameliorating GVHD post-HSCT

**DOI:** 10.1101/2025.01.16.633453

**Authors:** Duneia McManus, Sabrina N. Copsel, Brent J. Pffeifer, Dietlinde Wolf, Henry Barreras, Symon Ma, Ali Khodor, Seitaro Komai, Marina Burgos da Silva, Hajar Hazime, Miguel Gallardo, Marcel RM van den Brink, Maria T. Abreu, Geoffrey R. Hill, Victor L. Perez, Robert B. Levy

## Abstract

The current approach to minimize transplant-associated complications, including graft-versus-host disease (GVHD) includes long-term pharmacological immune suppression frequently accompanied by unwanted side effects. Advances in targeted immunotherapies regulating alloantigen responses in the recipient continue to reduce the need for pan-immunosuppression. Here, in vivo targeting of the TNF superfamily receptor 25 (TNFRSF25) and the high affinity IL-2 receptor with a TL1A-Ig fusion protein and low dose IL-2, respectively, was used to pretreat recipient mice prior to allogeneic-HSCT (aHSCT). Pretreatment induced Treg expansion persisting early post-aHSCT leading to diminished GVHD and improved transplant outcomes. Expansion was accompanied by an increase in frequency of stable and functionally active Tregs as evidenced by *in vitro* assays using cells from major GVHD target tissues including colon, liver, and eye. Importantly, pretreatment supported epithelial cell function/integrity, a diverse microbiome including reduction of pathologic bacteria overgrowth and promotion of butyrate producing bacteria, while maintaining physiologic levels of obligate/facultative anaerobes. Notably, using a sphingosine 1-phosphate receptor agonist to sequester T cells in lymphoid tissues, we found that the increased tissue Treg frequency included resident CD69^+^CD103^+^FoxP3^+^ hepatic Tregs. In contrast to infusion of donor Treg cells, the strategy developed here resulted in the presence of immunosuppressive target tissue environments in the recipient prior to the receipt of donor allo-reactive T cells and successful perseveration of GVL responses. We posit strategies that circumvent the need of producing large numbers of *ex-vivo* manipulated Tregs, may be accomplished through *in vivo* recipient Treg expansion, providing translational approaches to improve aHSCT outcomes.

**Graphical Abstract:** 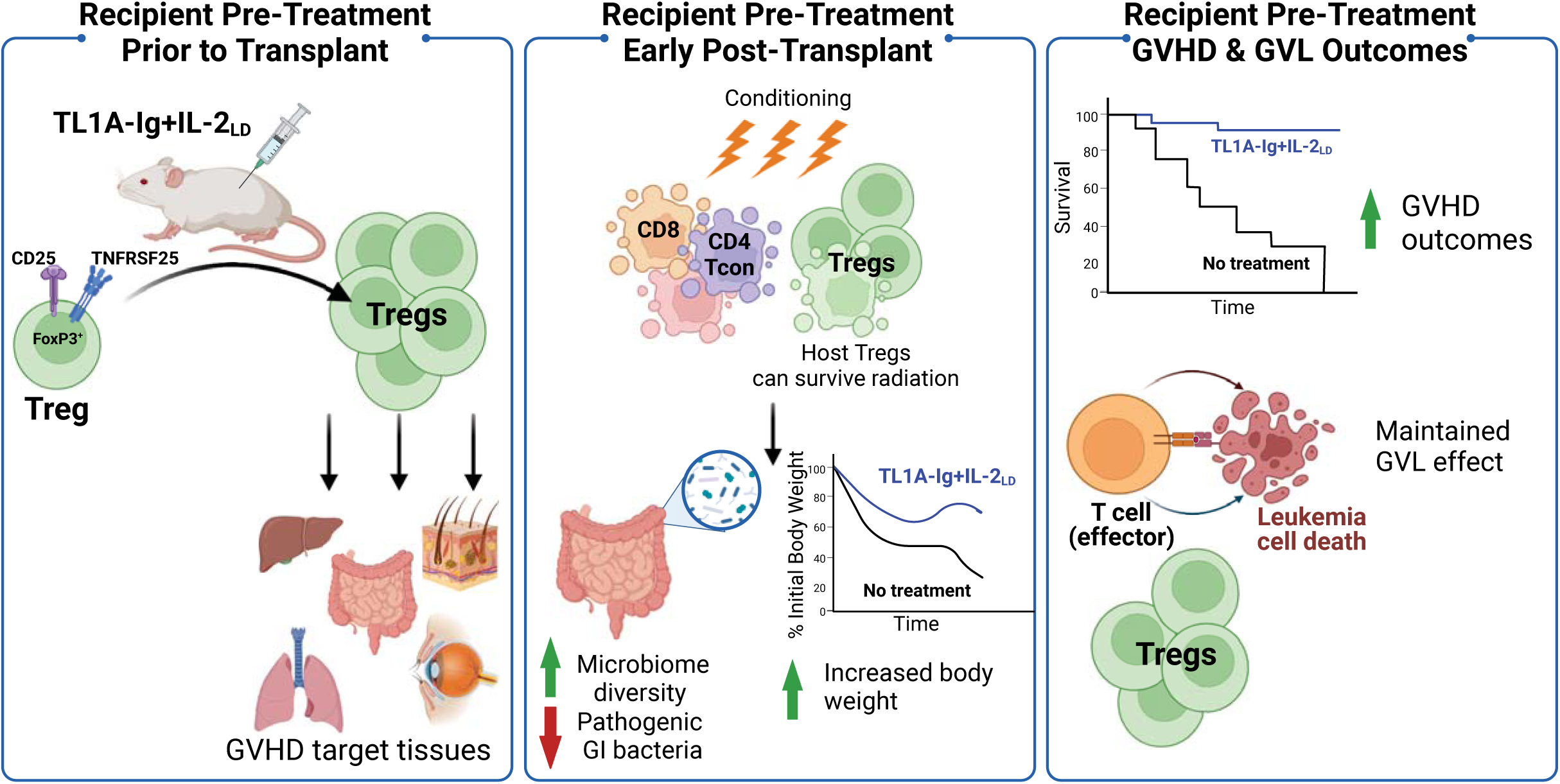

## Introduction

Allogeneic hematopoietic stem cell transplantation (aHSCT) is a curative procedure for patient with non-malignant and malignant disorders.^1^ However, one serious complication of aHSCT is GVHD, a donor T cell-mediated inflammatory process predominantly targeting the skin, gastro-intestinal tract, and liver.^2–4^ Clinically, GVHD prevention and treatment consists of pharmacological regimens which are pan-immunosuppressive placing patients at an increased risk for disease relapse or infectious complications. Additionally, many immunosuppressants have off-target effects causing other organ toxicities.^5,6^ As a potentially life-threatening complication, GVHD must be suppressed. However, maintaining graft-versus-leukemia (GVL) activity is critical to prevent tumor relapse.

Newer and alternative strategies to treat GVHD while preserving GVL have been the focus of current experimental and clinical research efforts. Several recently approved FDA compounds including kinase inhibitors and T cell activation regulators selectively target immune cells and show reduced drug toxicities.^7–14^ Immunotherapy is another strategy that may induce tolerance by manipulating regulatory immune cell populations. One such population shown to ameliorate GVHD is regulatory T cells (Tregs). Tregs, shown in experimental and clinical studies, can ameliorate GVHD across complete MHC mismatched aHSCT animal models and improve clinical symptoms of chronic GVHD in patients.^15,16^ To date, Treg based immunotherapeutic approaches for GVHD have primarily focused on the use of donor derived Tregs.^15,17–21^ While trials of adoptive Treg therapy in both preclinical and clinical aHSCT demonstrate safety and efficacy, there are significant challenges using adoptive Treg therapy, such as generating sufficient Treg numbers and potentially reduced stability. This may limit their use in individual transplants and off-the-shelf applicability.^15,18,22–29^

Notably, Tregs are more radio-/chemo-resistant than conventional CD4^+^ and CD8^+^ T cells and this likely contributes to a relative Treg enrichment early post-aHSCT.^30–36^ Recent studies show patient derived recipient T cells, including Tregs, residing in GVHD target tissues survive conditioning.^37,38^ These T cells were able to proliferate and were functionally competent >1 year post-aHSCT.^37,38^ We previously reported that recipient Tregs constitute the predominant component of the CD4^+^FoxP3^+^ compartment in lymph nodes for a number of weeks after radio-conditioning and auto-HSCT. Importantly these Tregs exhibit suppressive function.^39^ Therefore, we propose that *in vivo* expansion of Tregs in recipients prior to conditioning and aHSCT may be a useful strategy to diminish GVHD in target tissues and ameliorate overall disease. Here we report a new strategy where aHSCT recipients treated with compounds targeting and stimulating TNFRSF25 and the high-affinity IL-2 receptor only prior to transplantation expanding Tregs. This elevated recipient Treg frequency persisted for several weeks post-aHSCT, particularly in non-hematopoietic target tissues. The elevated Treg/T conv ratio was associated with suppressive environments as demonstrated by *in vitro* analyses and significant survival with diminished clinical GVHD. Importantly, GVL was maintained in recipients with greater Treg levels using an AML model. In total, these findings support the notion that pre-transplant manipulation of the recipient Treg compartment may be an effective therapeutic approach for GVHD prophylaxis while maintaining GVL.

## Methods

### Mice

BALB/c (H2^d^) mice were purchased from Taconic or Jackson Laboratory. C57BL/6J, B6-CD45.1, B6-FoxP3^RFP^, C3H.SW (H2^b^) and BALB/c-FoxP3^DTR^ (H2^d^) mice were bred in our facility. BALB/c^TNFRSF25-/-^ (H2^d^) were provided by Dr. H.Park (NCI, NIH).^40^ Mice (6 to 12 weeks old) were maintained in pathogen-free conditions at the University of Miami (UM) animal facilities. All animal use procedures were approved by the UM IAACUC.

### Flow cytometry and fluorescence activated cell sorting (FACS)

All antibodies used are listed in **Supplemental Table 1**. Single-cell suspensions were prepared from selected tissues/organs. Peripheral blood (PB) was collected in heparinized tubes and PBMCs were isolated using Ficoll-Paque^®^ (GE Healthcare, Chicago, IL). Cell surface and intracellular antibody staining was performed as previously described.^41^ Samples were run using either the LSR-Fortessa-HTS instrument (BD Biosciences, San Jose, CA) or a Cytek-Aurora (Cytek^®^ Biosciences, Bethesda, MD). Analysis was performed with FlowJo software (v.10.4.1, FlowJo, Ashland, OR). FCS files from aHSCT patients with and without GVHD underwent terraFlow multi-step pipeline unbiased analysis for summarization of key differences between groups.^42^

### Allogeneic HSCT and GVL

aHSCT was performed with sex and age-matched mice using either a major MHC-mismatch (B6→BALB/c) or a minor MHC-mismatch (C3H.SW→B6) model. Recipient mice received TBI conditioning: Day-1 (7.5 Gy) for major or Day0 (10.5 Gy) for minor MHC-mismatch. Transplantation was performed on Day0 for both models with T cell–depleted (TCD) bone marrow (BM), as described previously.^15^ Major MHC-mismatch recipients received (5 .5× 10^6^) TCD-BM with or without (5.5 × 10^5^) splenic T cells and minor MHC-mismatch recipients received (7× 10^6^) TCD-BM with or without (2.0 × 10^6^) splenic CD8^+^ T cells iv. Each recipient was monitored 3x/week for GVHD assessing overall survival, total body weight, and clinical score.^15^

For GVL assessment, BALB/c mice received 5x10^3^ BALB/c-MLL-AF9^GFP^ at the time of aHSCT (Day0) examining tumor burden by GFP^+^ cell frequency from blood and selected tissues on day of sacrifice using flow cytometry.

### TL1A-Ig+IL-2_LD_ *in vivo* administration

TL1A-Ig was generated as previously described.^43^ Our Treg expansion protocol consists of TL1A-Ig (40μg/mouse in PBS) given intraperitoneally (ip) daily on Days1-4 and recombinant human IL-2 (IL-2; 10,000 Units/mouse in PBS) given ip daily on Days4-6 (Roche, Indianapolis, IN).

### *In vivo* FTY720 administration

Mice were injected i.p. daily with 20 μg of FTY720 (Cat#SML0700-Sigma Aldrich, St Louis, MO) diluted in diH_2_O for the duration of the expansion protocol, Days1-6. Desired tissues were harvested and stained for flow cytometry on Day8.

### *In vitro* suppression assay

CD4^+^ cells were enriched from liver and lung cell isolates using the Easy Sep™ PE Positive Selection Kit II (StemCell Technologies-Cat#17684, Vancour, BC) and CD4-PE). Unfractionated single-cell suspensions of conjunctiva and lacrimal gland were used in some experiments. Cells were cultured in 96-well plates and activated with 1μg soluble anti-CD3 mAb (Clone-2C11). Cell cultures were imaged using an inverted microscope (Carl Zeiss Microscopy, White Plains, NY) obtaining brightfield images at 40X using a Keyence BZ-X710 Microscope (Itasca, IL) and counted by Trypan blue exclusion using Countess™ 3 Automated Cell Counter (Thermo-Fisher Scientific, Waltham, MA) after 120 hours.

### Spatial biology analysis using CODEX

Colons were harvested from mice that received a complete MHC mismatched aHSCT on Day+24; one group received TCD-BM only and another group received TCD-BM plus 0.5 x 10^6^ splenic T cells. Colons were flushed of stool using cold PBS. A 0.5 cm segment of the transverse colon was cut and placed perpendicularly into cryostat cassettes for OCT embedding and flash frozen. The tissue was sectioned transversely at 7μm and processed per Akoya’s Pre-Antibody Staining, Tissue Staining, and Post-Staining procedures using barcoded antibodies and corresponding barcode reporters listed in **Supplemental Table 2**. The Pheno-Cycler-Fusion software and instrument performed iterative, whole-slide imaging creating a high resolution QPTIFF file. This was later used in the QuPath (v0.5.1) environment to generate single cell annotations determining cell nuclei and cell boundaries. The generated single cell annotations and ultra-high parameter fluorescent data was exported to a csv file and used in R (Big Sur Intel build (8462)), RStudio (v2024.4.2) and the Seurat package (v5.0.3) to determine associated single cell phenotypic markers, dimensionality reduction and data visualizations.

### Microbiome Analysis

Stool samples from mice after aHSCT were taken on Day-8, -1, 0, +1, + 7, +14, +21, and +28 from MHC-mismatched recipients. DNA from stool was extracted and purified by bead-beating in phenol-chloroform as previously described.^44–46^ Genomic 16S ribosomal RNA V4/V5 regions were amplified by PCR and sequenced on the Illumina platform. Sequences were mapped and assigned taxonomically, as previously described.^44–46^ The microbial classification of predicted bacterial oxygen metabolism was adapted from Magnúsdóttir, S. *et al*.^47^ Bacteria were classified at the genus level when all species within that genus belonged to the same oxygen metabolism group **(Supplemental Table 3)** and classified at the species level when diverse metabolic groups were found within the same genus. A list of predicted butyrate producers was adapted from Haak, B. W. *et al.*.^48^

### Statistical Analysis

The number of animals/group are described in the legends. All non-spatial biology graphing and statistical analyses were performed using GraphPad Prism (San Diego, CA). Values shown in graphs represent the mean of each group ± SEM. Survival data were analyzed with the Mantel-Cox log-rank test. Nonparametric unpaired 2-tailed *t*-test was used for comparisons between 2 experimental groups. One-way ANOVA with Tukey’s multiple-comparison test was used between 3 or more experimental groups. A *p-*value less than .05 was considered significant and *p < 0.05, **p < 0.01, ***p < 0.001, ****p < 0.0001.

## Results

### Expansion of Tregs prior to conditioning alters CD4^+^FoxP3^+^ frequency post-TBI in the colon and peripheral lymphoid compartments

As previously reported, Treg frequency is diminished in the blood of patients with chronic GVHD **(Supplemental Figure 1A)**.^16^ We found Treg frequency differences in colonic tissues of mice Day+24 post-aHSCT using spatial biological analysis. Colon mononuclear cell assessment indicated significantly elevated levels of lymphocytes and macrophages and decreased levels of Tregs in BALB/c mice transplanted with BM+T, undergoing GVHD, vs. BM only **(Figure 1A&B, Supplemental Figure 1B-D).** This was accompanied by a significant reduction of Treg proliferation determined by decreased Ki67 expression **(Figure 1C)**.

**Figure 1.**
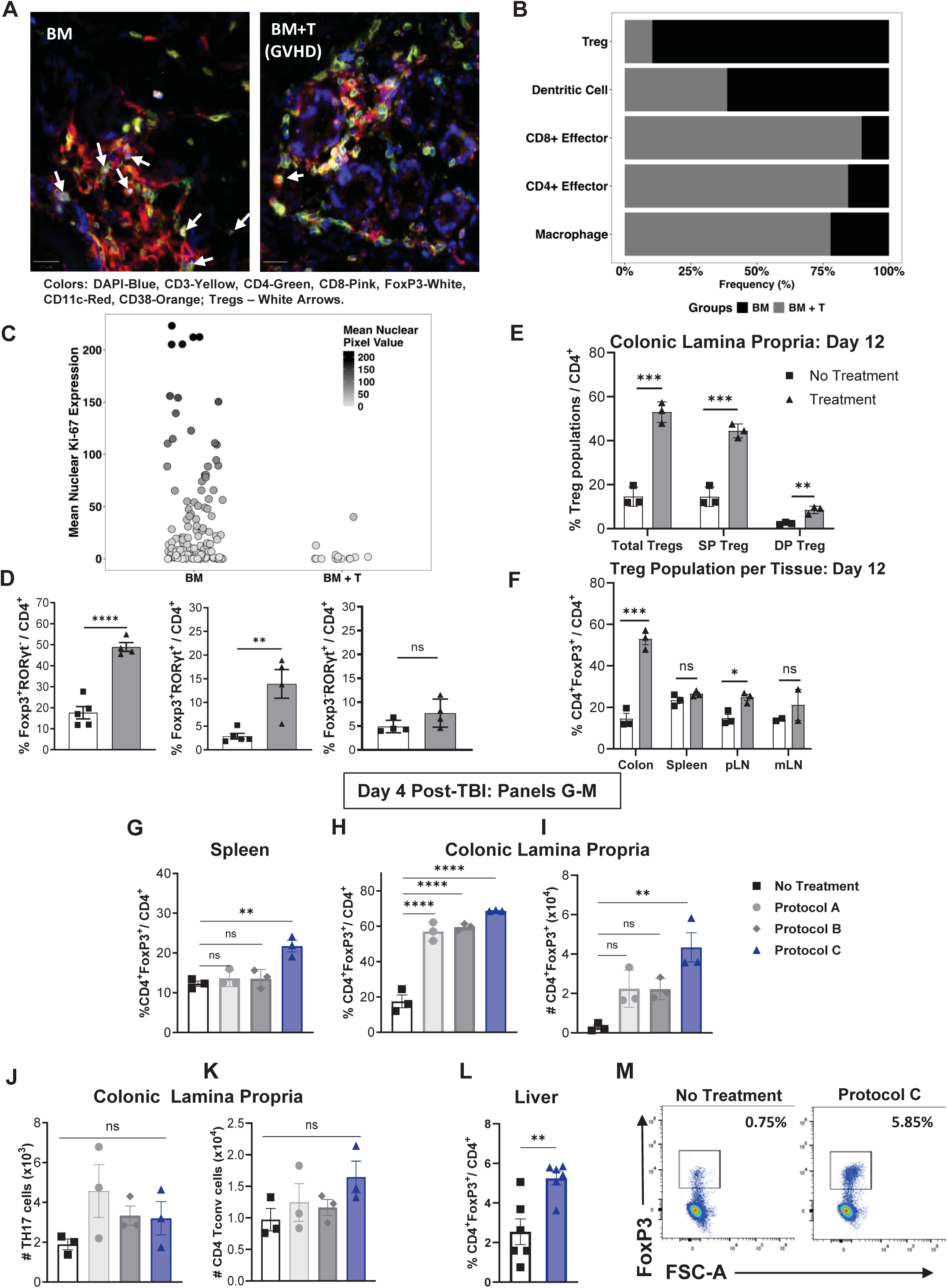
Optimizing *in vivo* Treg expansion protocol with TL1A-Ig+IL2_LD_ for Treg survival post-radiation. **(A)** QPTiff CODEX images illustrating a subset of cell populations (7-selected markers) highlighting diminished Treg numbers in tissue from BALB/c mice transplanted with BM+T undergoing GVHD vs. BM transplanted alone; white arrows show Treg locations in the images. **(B)** Select immune cell frequency separated by non-GVHD (BM only – Black bars) and GVHD (BM+T – Grey bars) from the whole colonic tissue, n = 2/group. **(C)** Mean nuclear Ki67 expression of Tregs separated by groups. Mice without GVHD (BM alone) contained a greater number of Tregs compared to GVHD animals (BM+T) with approximately 45% (compared to 0% in mice without GVHD) having moderate to high Ki67 expression indicating proliferation. **(D)** BALB/c mice were treated with TL1A-Ig+IL-2_LD_. Treg expansion in colonic lamina propria (LP) at day 7 but no detectable increase in colonic TH17 CD4^+^ T cells is shown. **(E & F)** BALB/c mice were assessed on Day 12, six days after completing TL1A-Ig+IL-2_LD_ treatment. **(E)** The colon LP shows a persistence in elevated total Tregs as well as Treg subsets (SP: CD4^+^FoxP3^+^RORγt^-^; DP: CD4^+^FoxP3^+^RORγt^+^) at day 12, and **(F)** this persistence is greatest in the colon vs other tissues. **(G-M)** BALB/c mice were treated with TL1A-Ig+IL-2_LD_ using different protocols **(Sup Figure 2C)** and received total body irradiation (TBI) (8.5 Gy) on Day -1. At Day 4 immune cells were assessed for Treg frequencies in the spleen **(G)** and colonic LP **(H).** Protocol C demonstrated significantly higher Treg frequencies and numbers compared to Protocols A, B, and No treatment **(G, H & I)**. Protocol C also shows no difference in Th17 **(J)** and CD4^+^ Tconv **(K)** numbers of the colon LP. **(L & M)** The liver exhibited a greater Treg frequency **(L)** using Protocol C. **(M)** shows representative flow cytometry plots gating on CD4^+^FoxP3^+^Treg cells using cell isolates from the liver. Data represent the mean ±SEM with *p<0.05; **p<0.01; ****p<0.0001 defining significance levels.

Administration of TL1A-Ig+IL-2_LD_ induces a strong expansion of Tregs in the blood, spleen, and LN of normal mice.^15,49^ Colonic lamina propria (LP) of mice receiving TL1A-Ig+IL-2_LD_ exhibited a strong elevation of CD4^+^FoxP3^+^RORγt^-^ and CD4^+^FoxP3^+^RORγt^+^ frequency, but virtually no change in CD4^+^FoxP3^-^RORγt^+^ (TH17) cells **(Figure 1D, Supplemental Figure 1E)**. The increased frequency of Tregs in the colonic LP **(Figure 1E)** remained elevated for 12 days post-treatment which was found to persist for a longer duration compared to Treg levels in the spleen and LN **(Figure 1F)**.

Recipient Tregs can persist post-conditioning (8.5 and 11 Gy) as shown in **(Supplemental Figure 2A-B)** and also following transplantation.^37,39,50,51^ Five days after TBI (8.5 Gy) Tregs were assessed in BALB/c mice using several protocols of expansion with TL1A-Ig+IL-2_LD_ **(Supplemental Figure 2C-F)**, and “Protocol C” resulted in the greatest percentage and numbers of Tregs in the PB, spleen, LP, and liver and was used in all subsequent experiments **(Figure 1G-I,L,M)**. Notably, neither CD4 T conventional nor TH17 total cell numbers were significantly altered following Treg expansion using Protocol C **(Figure 1J,K)**.

### Recipient TNFRSF25 and CD25 receptor stimulation prior to HSCT induces expansion of resident Tregs and a functionally suppressive milieu in GVHD target tissue

Following stimulation with TL1A-Ig+IL-2_LD_, there was marked expansion of Tregs (**Figure 2A**) and particularly the Treg Helios^+^ population **(Supplemental Figure 3A-B)** in multiple GVHD target tissues including the skin, liver, lung, and ocular tissue compared untreated.

**Figure 2.**
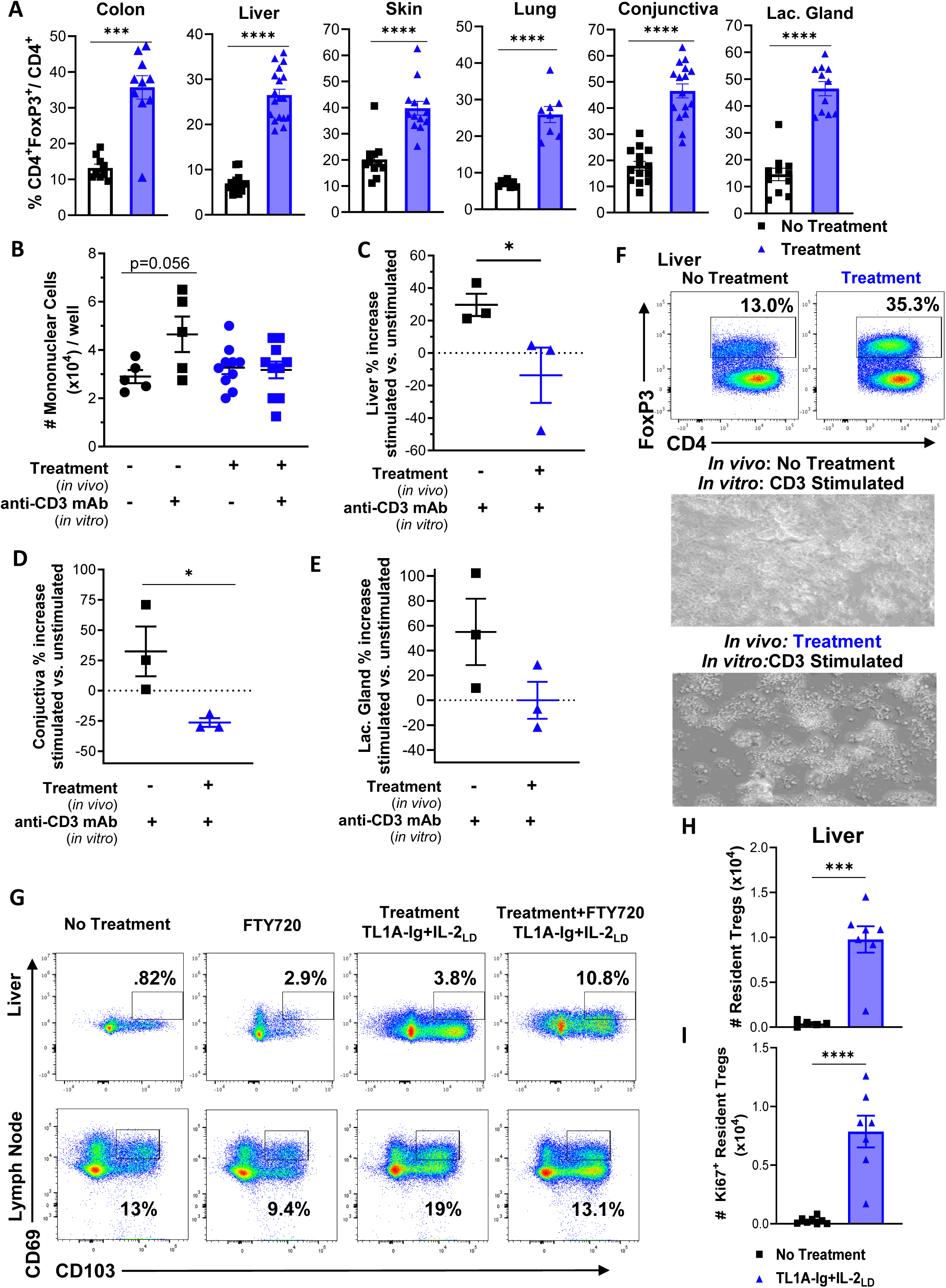
Recipient TNFRSF25 and CD25 receptor stimulation prior to HSCT induces expansion of tissue resident Tregs and a functionally suppressive environment in GVHD target tissues. **(A-F)** Assessment of GVHD target tissues of recipient BALB/c mice either untreated or treated with Protocol C (TL1A-Ig+IL-2_LD_) **(A)** Treg frequencies in non-hematopoietic tissue compartments -Colon, liver, skin, lung and ocular adnexa (conjunctiva and lacrimal gland) – on Day0 are shown. All GVHD target tissues show significant Treg expansion. **(B-E)** Measurement of suppressive capabilities of Treg cells isolated from GVHD target tissues on Day0 assessing *in vitro* proliferation using anti-CD3 mAb. **(C)** Cell number of colon lamina propria cultures were counted at hour 72 from Protocol C treated/untreated mice. Untreated mice showed higher cell counts compared to Protocol C treated mice after *in vitro* anti-CD3 stimulation. **(C-E)** *In vitro* studies assessing liver and the ocular adnexa. Each point represents 1 experiment, and each experiment had a minimum of 3 replicate wells for each group. On Day0, total lymphocytes from the liver were subjected to CD4^+^ enrichment via Easy Sep beads. These enriched CD4^+^ cells were then plated with anti-CD3 mAb to stimulate proliferation. **(C)** Percent increase of enriched liver mononuclear cells at hour 120 shows no net proliferation in cell counts when mice were treated with Protocol C. **(D)** Percent increase of total conjunctiva mononuclear cells at hour 120 if no treatment was given. **(E)** Percent increase of total lacrimal gland cells at hour 120 if no TL1A-Ig+IL-2_LD_ treatment was given. **(F)** Representative flow plots depicting frequency of FoxP3^+^ Tregs in wells at start of the cultures, and below representative images (40X magnification) of well health at hour 120 between *in vivo* treated or untreated, respectively. **(G - I)** The S1PR modulator, FTY720, was used to delineate whether the TL1A-Ig+IL-2_LD_ expanded Tregs traffic from hematopoietic sites or proliferate in non-hematopoietic GVHD target tissues. **(G)** Gating strategy for resident FoxP3^+^ Tregs (CD69^+^CD103^+^) from liver (upper) and cervical lymph nodes (lower panel). **(H)** Absolute numbers of resident (CD69^+^CD103^+^) Tregs in liver are significantly increased after TL1A-Ig+IL-2_LD_ treatment. **(I)** The vast majority of Tregs highly express the proliferation marker, Ki67, indicating robust tissue resident Treg expansion. Data represent the mean ±SEM with *p<0.05; **p<0.01; ****p<0.0001 defining significance levels.

To assess the functional ability of Tregs from non-hematopoietic tissues, lymphocytes from the colon, liver, and ocular adnexa (conjunctiva and lacrimal gland) from untreated and treated mice were examined *in vitro*. Following anti-CD3mAb stimulation, tissues with expanded Tregs exhibited a significantly lower proliferation rate **(Figure 2B-F)**, suggesting that these tissue milieus reflect a functionally suppressive environment.

To address if tissue resident (TR) Tregs were targeted by TL1A-Ig+IL-2_LD_, CD69^+^CD103^+^FoxP3^+^ cells were examined using FTY720 to sequester non-resident T cells to the LNs **(Supplemental Figure 3C)**. Notably, TL1A-Ig+IL-2_LD_ treatment in combination with FTY720, resulted in a significant increase of Ki67^+^ TR Tregs in the liver at time of sacrifice (**Figure 2G-I**, **Supplemental Figure 3D-E)**. These results support the notion that TL1A-Ig+IL-2_LD_ administration stimulates resident Treg expansion within non-hematopoietic tissues.

### Administration of TNFRSF25 and/or CD25 agonists prior to conditioning results in elevated colonic Treg frequency post-transplant accompanied by improved GVHD outcome

To examine if pre-HSCT Treg expansion in recipients alters the outcome of aHSCT regarding GVHD, TL1A-Ig+IL-2_LD_ pretreated mice were transplanted across a complete donor/recipient MHC disparity **(Supplemental Figure 4A)**. Pretreated mice exhibited a marked increase (Day-2) in the frequency of PB Tregs **(Figure 3A)**.

**Figure 3.**
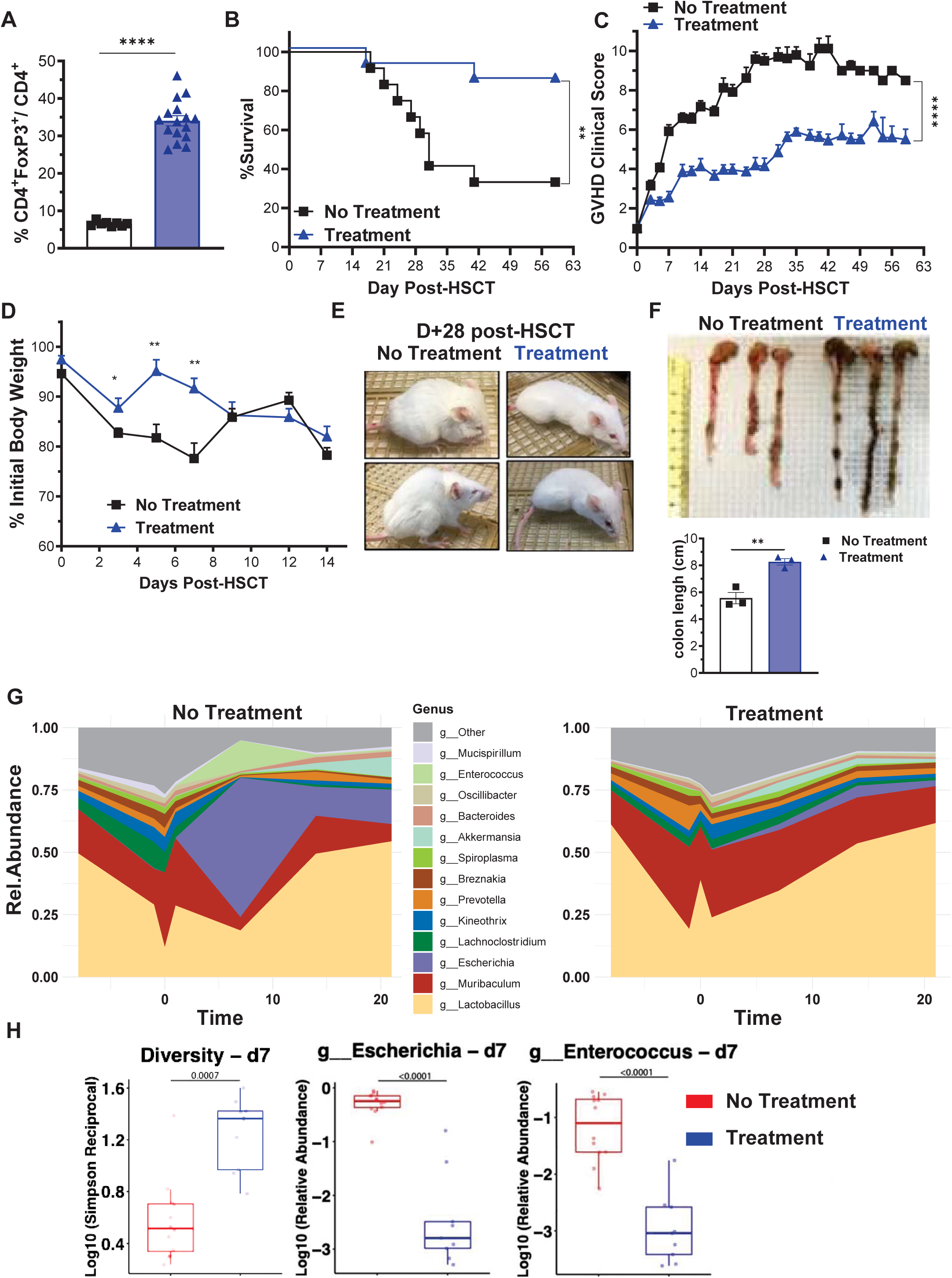
Recipient Treg expansion with TL1A-Ig+IL-2_LD_ prior to transplant significantly diminished GVHD severity, promoted colonic biome diversity, and improved overall survival. **(A-F)** BALB/c mice were treated with TL1A-Ig+IL-2_LD_ prior to TBI. aHSCT was performed using a B6 (donor) BALB/c (recipient) major MHC-mismatch model. CD45.1 B6 (H2^b^) T cell depleted (TCD) bone marrow (BM) (5.5x10^6^) and an adjusted spleen cell number containing 0.55x10^6^ T cells were transplanted into BALB/c (H2^d^) recipients. TBI was 7.25 Gy given on Day -1. The presented aHSCT data was pooled from 2 independent experiments representing 16 mice/group. **(A)** Confirmation of Treg expansion (CD4^+^FoxP3^+^/CD4^+^) in peripheral blood on Day -2. **(B-F)** Panels show survival **(B)**, GVHD clinical score **(C)**, percent weight loss **(D)**, general mouse appearance **(E)**, and colon lengths **(F)**. **(B)** Overall survival was 38% without recipient TL1A-Ig+IL-2_LD_ treatment versus 89% with recipient TL1A-Ig+IL-2_LD_ treatment. There was a significant difference in GVHD clinical scores over 63 days **(C)** and less percent weight loss early after aHSCT in TL1A-Ig+IL-2_LD_ treated mice**(D). (E)** Representative photographs depict animal appearance at Day +28 post-aHSCT. **(F)** Colon lengths of mice (n=6, 3 from each group) at Day +42 post aHSCT. **(G)** Relative abundance of bacterial microbiome over time in untreated and TL1A-Ig+IL-2_LD_ pretreated recipients. Stool was assessed at Day 0, Day +7, Day +10, Day +14, and Day +20. **(H)** TL1A-Ig+IL-2_LD_ treated recipients had greater microbiome diversity D+7. Bacterial diversity (Simpson Reciprocal) and specific genus-level relative abundance of *Escherichia* and *Enterococcus* Data represent the mean ±SEM with *p<0.05; **p<0.01; ****p<0.0001 defining significance levels.

Following recipient Treg expansion in BALB/c (H2^d^) mice prior to TBI and HSCT with B6 (H2^b^) TCD-BM ± splenocytes, recipients receiving pretreatment had increased survival and diminished GVHD clinical scores **(Figure 3B,C, Supplemental Figure 4B)**. Initial weight loss early post-HSCT was reduced in treated animals and overall appearance was clearly improved **(Figure 3D,E)**. Colon integrity determined by FITC-dextran leakage **(Supplemental Figure 4C)**, and colon lengths were greater in treated compared with untreated recipients six weeks post-transplant **(Figure 3F)**.

The intestinal microbiome has been found to become less heterogenous in GVHD, thus fecal material was collected and analyzed for bacterial species and relative diversity.^52–54^ Recipient pretreatment resulted in a more diverse intestinal microbiome following aHSCT compared to untreated BALB/c recipients **(Figure 3G,H).** Moreover, in the absence of pretreatment, higher levels of pathogenic species (ex. Escherichia and Enterococcus) were identified **(Figure 3H)**. Maintenance of butyrate producing and obligate anaerobe microbes, as well as epithelial H_2_O_2_ production were observed in pre-treated compared to untreated recipients **(Supplemental Figure4D-F),** consistent with a more physiologic GI environment.

To evaluate the effect on GVHD outcomes of individual TNFRSF25 or CD25 compared to combinatorial stimulation, mice were administered only TL1A-Ig, only IL-2_LD,_ or the combination to augment the Treg compartment (**Figure 4A,B**, **Supplemental Figure5A-C)**. Post-aHSCT, outcomes assessed by overall survival and GVHD clinical scores indicated that pretreatment with IL-2_LD_ alone had minimal Treg effects whereas pretreatments which included TL1A-Ig resulted in reduced colonic cellular infiltrates, significant improvement in survival and clinical scores compared with untreated recipients **(Figure 4A-C**).

**Figure 4.**
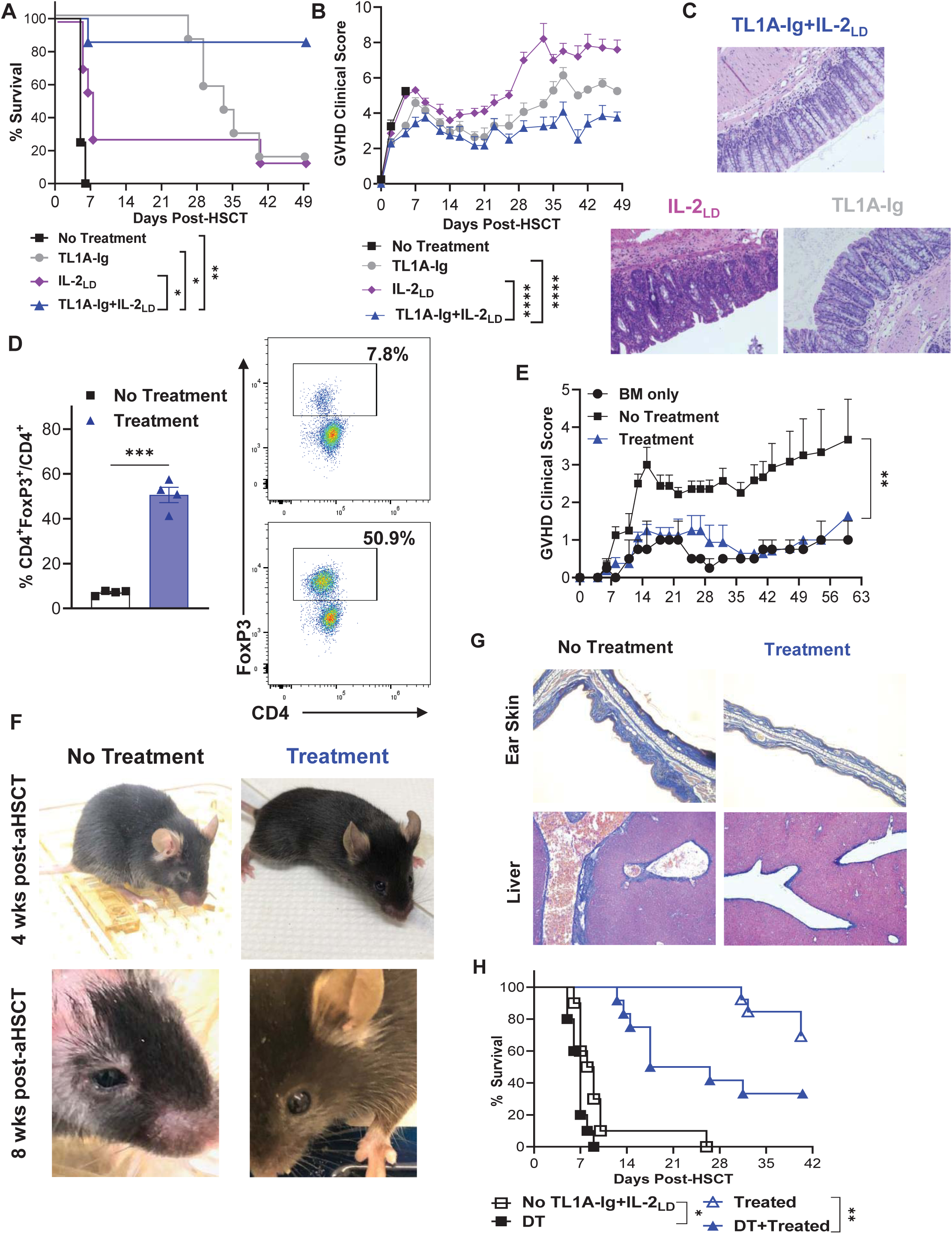
Combinatorial vs individual TL1A-Ig and IL-2 treatment prior to major and minor-MHC-mismatched aHSCT. **(A-C)** Evaluation of recipients after individual TNFRSF25 or high affinity IL-2 receptor stimulation compared to combined stimulation prior to conditioning and aHSCT. Mice were administered TL1A-Ig only, IL-2_LD_ only, or TL1A-Ig+IL-2_LD_ (see Supplemental Figure 4A for details). Allogeneic HSCT was performed using a B6→BALB/c donor/recipient mouse model. B6-FoxP3^RFP^ (H2^b^) TCD-BM (5.5x10^6^) cells and an adjusted spleen cell number containing 6.5x10^5^ T cells were transplanted into BALB/c (H2^d^) recipients after TBI conditioning (7.75 Gy) on Day-1. **(A)** Overall survival, **(B)** GVHD clinical score and **(C)** representative colon histological images of tissue harvested at Day+56. Combinatorial pretreatment resulted in the best outcomes with fewer lymphocytic infiltrates in the colonic tissue. **(D-G)** The combinatorial treatment was examined in a second (low lethality) aHSCT model. MHC-matched, minor antigen mismatched C3H.SW (H2^b^) TCD-BM (7x10^6^) cells and 2x10^6^ adjusted splenic CD8^+^ T cells were transplanted into B6 (H2^b^) recipients after TBI conditioning (10.5 Gy) on Day0. **(D)** At Day-2, mice Treg frequency was assessed showing significant CD4^+^FoxP3^+^/CD4^+^ expansion with pretreatment with representative flow plots. **(E)** GVHD clinical scores were less severe in pretreated recipients. **(F)** Representative images of mice show appearances at 4 and 8weeks post-aHSCT. Pretreated recipients exhibited less skin and ocular GVHD involvement. **(G)** Collagen deposition (Masson’s Trichrome, blue) in the skin and liver of mice 8 weeks post-aHSCT. In treated recipients, note the thin epidermis, scant dermis collagen staining and absence of inflammatory cells versus untreated recipients exhibiting a thickened epidermis, extensive collagen deposition in the dermis and a modest infiltration of chronic inflammatory cells, magnification =10X. Heightened collagen and hepatic peri-portal infiltrates were detected in untreated MHC-matched recipients. **(H)** Recipient Treg depletion immediately prior to transplant exacerbates GVHD related lethality. BALB/c diphtheria toxin receptor (DTR) mice were untreated or pretreated with TL1A-Ig+IL-2_LD_ (Day-7 to Day-2) prior to transplant with B6 TCD-BM± splenocytes. Diphtheria toxin (1ug ip) was administered on Day-1 prior to TBI. Depleting recipient Tregs reduced – but did not abolish – protection resulting from TL1A-Ig+IL-2_LD_ pretreatment.

Since combinatorial pretreatment resulted in the greatest protection from GVHD, the protocol was examined in a second (low lethality) aHSCT model. MHC-matched, minor antigen mismatched C3H.SW (H2^b^) BM + T cells were transplanted into B6 (H2^b^) recipients. As anticipated, Treg levels were markedly expanded with TL1A-Ig+IL-2_LD_ prior to HSCT (**Figure 4D**). Clinical scores were significantly diminished in treated recipients **(Figure 4E)**. Representative photographs **(Figure 4F)** 4 and 8 weeks post-transplant illustrate the improved appearance of treated mice accompanied by less histopathological involvement in skin, and liver and less eye involvement **(Figure 4F,G).**

To assess if the depletion of expanded Tregs affected GVHD outcomes, FoxP3-diphtheria toxin (DT) receptor (BALB/c-FoxP3 ^DTR–eGFP^) mice were TL1A-Ig+IL-2_LD_ pretreated and administered DT immediately before TBI **(Supplemental Figure 5D-F)**. As anticipated, animals pretreated with TL1A-Ig+IL-2_LD_ exhibited significantly greater overall survival compared to untreated recipients. Removal of Tregs considerably decreased percent survival in TL1A-Ig+IL-2_LD_ treated animals, providing evidence that Tregs are important in the amelioration of GVHD (**Figure 4H**). Consistent with the importance of Tregs in GVHD suppression observed, removal of these cells *in vitro* resulted in elevated anti-CD3 mAb stimulated T cell responses **(Supplemental Figure 5G)**. This activity was TNFRSF25 dependent, as TNFRSF25 receptor knock-out mice did not exhibit significantly increased Tregs *in vivo* following TL1A-Ig+ IL-2_LD_ treatment or suppressive activity *in vitro* **(Supplemental Figure 5H,I)**.

### Administration of TL1A-Ig+IL-2_LD_ before conditioning and transplant resulted in a strong preponderance of recipient Tregs in the colonic LP and spleen post-HSCT

Administration of TL1A-Ig+IL-2_LD_ into normal, non-transplanted animals resulted in a strong increase in Treg levels in the colonic LP which persisted for 12 days **(Figure 1E)**. To investigate if the frequency was also elevated early after conditioning and transplant, Treg levels were examined in the colons of TL1A-Ig+IL-2_LD_ treated and untreated BALB/c recipients four days post-HSCT with MHC-mismatched donor B6 cells **(Figure 5A-E)**. Assessment of the colonic LP revealed that CD4^+^FoxP3^+^Treg frequency was significantly elevated in this compartment as initially observed in the absence of transplant **(Figure 1D-F).** Both single (FoxP3^+^RORγt^-^) and double (FoxP3^+^RORγt^+^) positive Treg populations exhibited elevated frequency and numbers (**Figure 5A,B**). In contrast, frequency of TH17 (CD4^+^FoxP3^-^RORγt^+^), total CD4, and CD8 cells did not exhibit any increase in these Treg expanded animals **(Figure 5A,C,D)**. The Treg frequency was also elevated in the spleen four days post-HSCT, however the relative increase compared to the colon was lower while the CD8 frequency was significantly reduced (**Figure 5F-H**). To corroborate selective Treg increase and persistence at this time in the colon, pretreatment with an anti-TNFRSF25 agonistic mAb (mPTX35) and IL-2_LD_ was administered to BALB/c mice, prior to B6 transplant **(Supplemental Figure 6A,B)**. As anticipated, a significant increase in single and double positive Tregs was observed.

**Figure 5:**
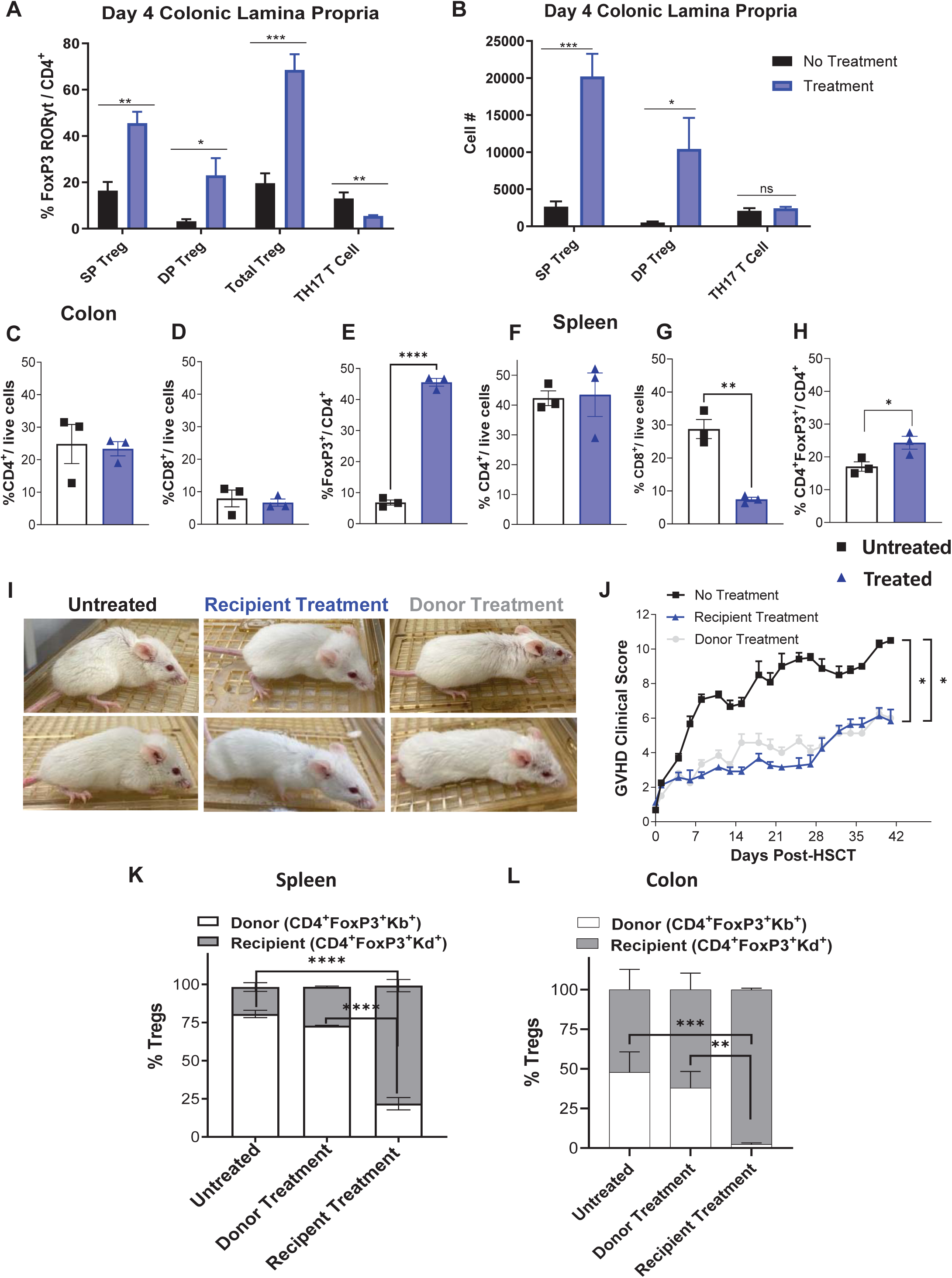
Donor and recipient expanded Tregs significantly diminished GVHD severity. **(A-H)** Recipient BALB/c mice were pretreated with TL1A-Ig+IL-2_LD_. An aHSCT (B6→BALB/c) was performed on Day 0 (n=3 mice/group) following conditioning on Day-1. Tissue resident colonic lamina propria (LP) lymphocytes were assessed at Day+4 post-aHSCT. **(A)** The frequencies and **(B)** absolute numbers of FoxP3^+^RORyt^-^ / CD4^+^ T cells, FoxP3^+^RORyt^+^ / CD4^+^ T cells, total Tregs, and TH17 / CD4^+^ T cells are shown. **(C, D, & E)** The frequencies of CD4^+^, CD8^+^, and FoxP3^+^ / CD4^+^ T cells from colon LP and **(F, G, & H)** spleen cells are shown. **(I, J)** Pre-treated BALB/c recipients of B6 aHSCT exhibited improved appearance (Day+28 post-aHSCT) and significantly improved clinical scores depicting reduced GVHD scores in both, donor and recipient TL1A-Ig+IL-2_LD_ treated groups compared to the untreated group. **(K,L)** TL1A-Ig+IL-2_LD_ pre-treated recipients demonstrated a preponderance of recipient Tregs in the colon and spleen. Donor (H2^b^) and recipient (H2^d^) Treg frequencies on Day+5 post-aHSCT (B6èBALB/c) in the spleen **(K)** and colon **(L)** were analyzed. TL1A-Ig+IL-2_LD_ pre-treatment (n=8 mice/group) resulted in a majority and almost exclusively Tregs of recipient origin in both, spleen and colon, respectively. Notably, at Day+5 post-aHSCT, in contrast to the spleen **(K)**, recipient Tregs in the colon (**L**) still represented the majority of the Treg population **(L)** after transplant with Treg expanded donor cells. Data represent the mean ±SEM with *p<0.05; **p<0.01; ****p<0.0001 defining significance levels.

The elevated frequency of Tregs in recipients post-transplant could be comprised of both donor and host Tregs. A transplant was performed with recipients either untreated, transplanted with cells from Treg expanded donors, or pretreated with TL1A-Ig+IL-2_LD_ **(Figure 5I,J)**.^15^ GVHD was markedly ameliorated in both latter groups **(Supplemental Figure 6C)**. In both, untreated recipients and recipients receiving Treg expanded donor spleen cells, the preponderance of Tregs in the spleen were donor derived 4-5 days post-aHSCT **(Figure 5K)**. In contrast, recipient pre-treatment established a marked host Treg predominance (**Figure 5K,L**). Notably, recipient Tregs comprised ∼50% in the colon of untreated or mice receiving transplants from Treg expanded donors at this time (**Figure 5L**). Strikingly, pretreatment resulted in almost complete recipient Treg presence (**Figure 5K,L**). Therefore, a preponderance of recipient Tregs was associated with improved transplant outcomes correlated with diminished proliferation of donor Tconv CD4 cells (**Supplemental Figure 6D,E**)

### The anti-leukemia response is maintained in TL1A-Ig+IL-2_LD_ treated recipients following aHSCT with MLL-AF9 leukemia cells

Groups of BALB/c pretreated recipients were transplanted with B6 TCD-BM ± splenocytes and BALB/c-MLL-AF9^GFP^ AML cells to determine if GVL responses were maintained in TL1A-Ig+IL-2_LD_ pretreated mice. Recipients of B6 BM only + MLL-AF9^GFP^ cells all died by 4 weeks post-aHSCT because of tumor outgrowth (**Figure 6A,B**). Untreated recipients transplanted with B6 TCD-BM+splenic T cells survived longer as a consequence of GVL (**Figure 6A,B**). As anticipated, TL1A-Ig+IL-2_LD_ treated recipients showed diminished clinical manifestations of GVHD 4 weeks post-aHSCT (**Figure 6C**). We identified MLL-AF9^GPF^ cells in BM, liver, reproductive organs (uterus and ovaries), and spinal cord of recipients transplanted with B6 BM alone beginning ∼4 weeks post-aHSCT; however, tumor cells were not detected in pretreated recipients receiving the identical transplant with TCD-BM+splenic T cells at week 6 post-aHSCT **(Figure 6D-G)**. These findings indicate that pretreatment of recipients using TNFRSF25 and CD25 agonists did not impair their ability to mount anti-leukemia cell responses following aHSCT.

**Figure 6.**
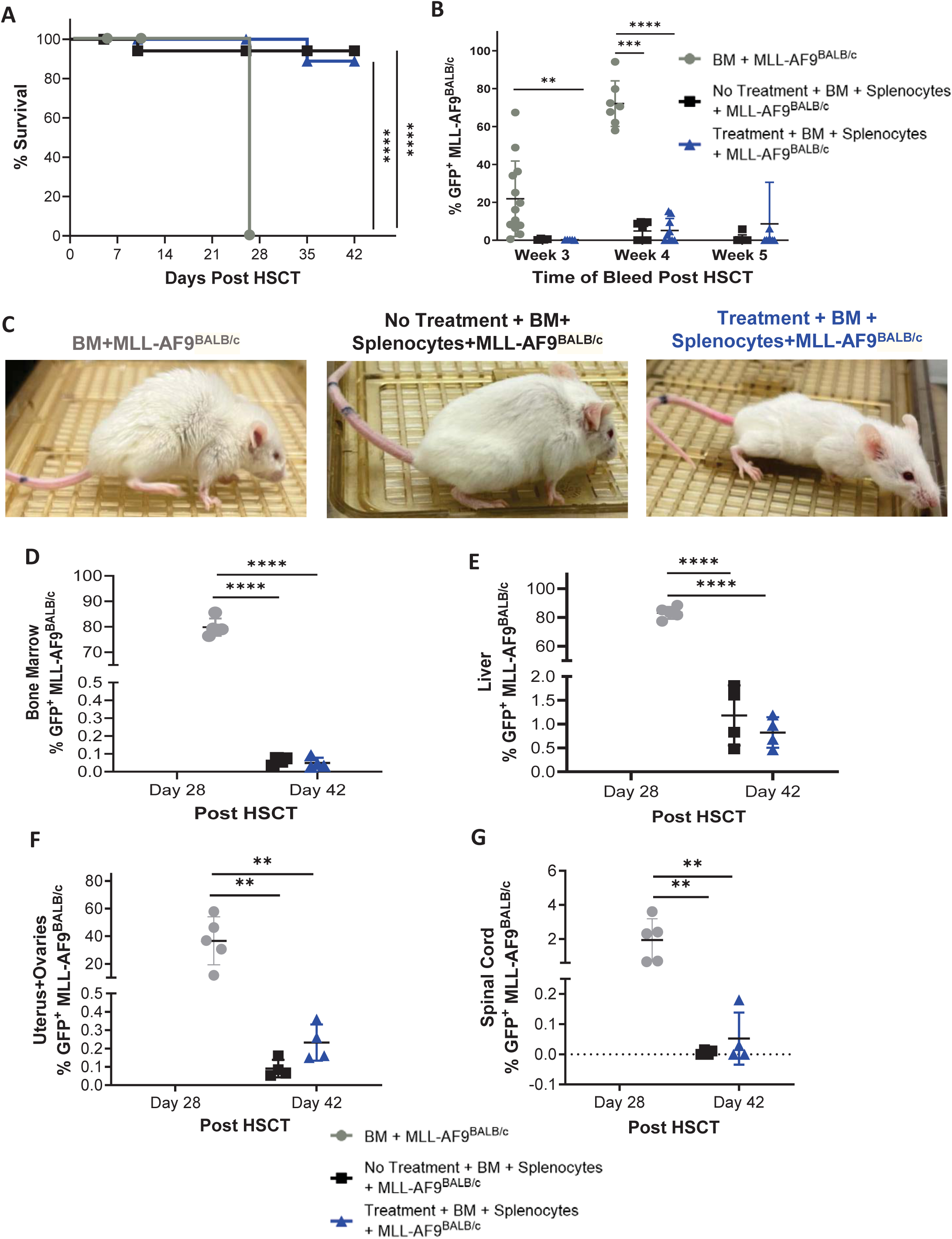
Recipient mice pretreated with TL1A-Ig+IL-2_LD_ prior to aHSCT enable GVL responses concomitant with GVHD amelioration. A B6 (donor) → BALB/c (recipient) aHSCT with and without recipient TL1A-Ig+IL-2_LD_ pretreatment was performed. All groups received 5x10^3^ BALB/c-MLL-AF9^GFP^ cells (iv) at the time of aHSCT, (n=9-13 mice/group). **(A)** Overall survival. No animals survived in the BM only group versus > 85% for the (BM + T) and (BM + T with recipient treatment) groups. **(B)** BALB/c-MLL-AF9^GFP^ cell frequency in peripheral blood at 3, 4, and 5 weeks post-aHSCT. Treg expansion in the recipient group enabled a GVL response against BALB/c-MLL-AF9^GFP^ cells that was as effective as the response in untreated recipients (BM+T) denoted by minimal detection of GFP expression. **(C)** Representative photographs of mice from each group at 4 weeks post aHSCT. **(D-G)** BALB/c-MLL-AF9^GFP^ frequency in hematopoietic - bone marrow **(D)** and non-hematopoietic compartments: liver **(E),** uterus and ovaries **(F)**, and spinal cord **(G)** at days 28 and 42 post-aHSCT.

## Discussion

Tregs can alter immune responses under physiologic conditions and their application as cellular therapy to diminish those responses in transplantation and autoimmune diseases continues to show promise.^55–61^ Expansion of Tregs using IL-2 muteins and other reagents to increase this compartment *in vivo* continues to advance.^56,62–64^ Our laboratory has developed strategies to augment Tregs *in vivo* by targeting the TNFRSF25 receptor alone and together with IL-2_LD_.^15,43,55,65^ While most Treg expansion approaches in HSCT have focused on the use of donor derived Tregs, the present investigation reports for the first time a translatable strategy to expand Tregs in recipients prior to transplant. Results demonstrated that elevation in CD4^+^FoxP3^+^ Treg frequency and numbers post-transplant was associated with diminished GVHD and improved outcomes following aHSCT across major and minor histocompatibility disparities.^18,19,26,66–70^

TNFRSF25 is a costimulatory molecule upregulated on CD4^+^ Tconv and CD8^+^ T cells following activation.^71,72^ In contrast, this molecule is constitutively expressed on Tregs and in normal mice, we reported its signaling can rapidly expand these cells after administering an agonistic fusion protein (TL1A-Ig) or mAb (4C12).^15,43,55,65^ We previously identified a six-day *in vivo* protocol resulting in marked Treg expansion in multiple compartments, including blood, LN, spleen, and colon but not in BM.^15^ A decrease in Treg frequency in the PB – as well as dysfunction - has been reported in patients with cGVHD.^16,73–75^ The present study corroborated this observation **(Supplemental Figure1A)** and also identified a deficit in Treg levels and function in the colon of mice with GVHD using spatial biology **(Figure 1A-C)**. We hypothesized expanding the Treg compartment in the recipient prior to transplant may prevent a Treg deficit. Attempts to expand the compartment immediately post-transplant with mAb targeting TNFRSF25 exacerbated GVHD due to stimulation of activated donor T cells.^76^ We therefore applied our Treg expansion protocol immediately prior to HSCT. The TL1A-Ig FP and not mAb was selected due to its short half-life to avoid potential stimulation of donor allo-reactive T cells and resulted in significantly improved transplant outcomes.^43^

Tissue resident (TR) Tregs with distinct transcriptional and epigenetic programs reside in nonlymphoid compartments including important target GVHD tissues.^77^ TL1A-Ig+IL-2_LD_ administration was shown to significantly elevate Tregs in the skin, liver, colon, eye, and lung. Notably, after blocking egress of Tregs from LN with FTY720, we found that CD103^+^CD69^+^ TR Tregs were markedly increased in the liver, indicating their expansion in the target tissues. Importantly, we detected a preponderance of recipient Tregs in multiple compartments during the first week post-transplant following pre-transplant expansion with TL1A-Ig+IL-2_LD_. Since we found that *in vitro* stimulated T cell responses from tissues were suppressed following Treg expansion, we posit that pretreatment promoted a locally suppressive environment persisting early post-transplant, thereby reducing GVHD. Consistent with this interpretation, deletion of recipient Tregs via diphtheria toxin injection following expansion one day prior to aHSCT resulted in significantly reduced but not abolished protection. Therefore, we speculate that other populations could also be contributing to the observed effect, and studies will investigate this possibility. Thus, while TL1A-Ig alone elevated Tregs in nonhematopoietic tissues comparable to combination treatment, GVHD outcomes were best following the latter treatment.

Recent reports in transplant patients have shown that recipient T cells - including Tregs - while absent in blood, were present in GVHD target tissues >1 year post-aHSCT.^37,38^ These findings indicated such cells survived conditioning and moreover, were functionally competent and able to proliferate. ^37,38^ Tregs are reportedly more radio and cyclophosphamide resistant than conventional CD4^+^ and CD8^+^ T cells, which may contribute to their relative enrichment post-aHSCT.^30,51,78^ Recipient Tregs comprised the predominant component of the CD4^+^FoxP3^+^ compartment in LNs five weeks after autologous-HSCT in mice, and these persistent Tregs were found to mediate suppressive function.^39,50^ Notably, recipient Tregs were present following TBI, cyclophosphamide, and busulfan conditioning protocols.^39^ Consistent with these observations, we found elevated levels of recipient Tregs in multiple tissues following their expansion pre-transplant. Helios has been reported to regulate and stabilize Foxp3 expression as well as Treg-suppressive function and survival, as Helios deficiency results in the loss of Treg lineage in inflammatory conditions.^79–84^ Notably, Helios’ expression was substantially increased in TL1A-Ig+IL-2_LD_ expanded Tregs in multiple tissues at the time of conditioning and transplant **(Supplemental Figure 3A,B)**.

Perturbations of the GI microbiome can cause immune dysregulation following aHSCT and major alterations including reduced diversity has been identified in the colon.^52–54,85–87^ Pre-treatment with TL1A-Ig+IL-2_LD_ maintained bacterial species diversity and promoted butyrate producing species while diminishing pathogenic bacterial overgrowth **(Supplemental Figure 4D)**. These modifications are of note as butyrate plays a key role in the promotion of Treg differentiation and stability, improves-suppressive function and enhances metabolic support and expansion in the gut.^88–90^ Hypoxia in the colon regulates the bacterial species present and in pretreated mice, there was a higher ratio of obligate to facultative anaerobic species consistent with a more homeostatic hypoxia status **(Supplemental Figure 4E)**. Duox2 signaling in the gut is critical for H_2_O_2_ production, important for microbial defense, and involved in regulating epithelial cell proliferation and repair.^91–94^ Commensal bacteria induces Duox2 expression under physiologic conditions while dysbiosis can lead to its dysregulation leading to epithelial injury and impairing epithelial repair mechanisms.^91–93,95,96^ While transplant conditioning here was associated with transient bacterial dysregulation and elevated H_2_O_2_ levels early post-aHSCT, animals with GVHD exhibited sustained, elevated levels of colonic H_2_O_2_ at least 3 weeks post-aHSCT. In contrast, H_2_O_2_ production in pre-treated recipients of BM+T cells was comparable to levels in recipients of BM alone **(Supplemental Figure 4F)**. Therefore, we conclude that TL1A-Ig+IL-2_LD_ administration promoted epithelial cell integrity and preservation of indigenous microbiota.

Maintaining GVL responses while ameliorating GVHD despite Treg expansion is important for translational applicability. MLL-AF9^GFP^ leukemia cells were transplanted together with BM±spleocytes to assess for anti-tumor activity. TL1A-Ig+IL-2_LD_ pretreated animals mediated GVL responses in the presence of elevated Treg levels as indicated by absence of leukemia cells in multiple compartments comparable to non-pretreated recipients of BM±splenocyte transplants. We have reported previously that TL1A-Ig+IL-2_LD_ does not increase Treg levels in the marrow, a key site of MLL-AF9 expansion.^15^ Pretreatment elevated Treg levels in GVHD target tissues i.e. liver, skin and eye. This differential tissue Treg expansion together with potential anti-tumor antigen-specific responses may explain why, as others have reported, Treg suppression of GVHD (alloantigen-specific T cells) here did not abolish anti-leukemia cell responses.^15,19^

In total, manipulating the recipient Treg compartment with TL1A-Ig+IL-2_LD_ expanded Treg cells in both hematopoietic and non-hematopoietic tissues, including the colon - central to the induction of GVHD.^55,97^ As a result of increasing their numbers pre-aHSCT followed by persistence post-aHSCT, this leads to elevation of recipient Tregs in GVHD target tissues several weeks post-transplant supporting the notion that the observed Treg deficit in GVHD can be reduced using this approach. In contrast to the infusion of donor Tregs, this pretreatment strategy we developed results in the presence of immune-suppressive target tissue environments in the recipient prior to donor allo-reactive T cell infusion. We posit strategies that circumvent the need of producing large numbers of Tregs ex-vivo by manipulating this compartment *in vivo*, may provide an effective therapeutic approach for GVHD prophylaxis while maintaining GVL.

## Acknowledgements

We want to thank Sylvester Cancer Center Flow Cytometry Shared Resource Core for their help and support. Special acknowledgement is extended to Core staff members specifically, Patricia Guevara, Shannon Saigh, PhD, Brit Chapman, and Eric Weider, PhD, Core Director for their help with terraFlow and all flow cytometric analysis.

The authors are highly appreciative of the help by Drs. Trent Wang, Eric Weider, and Cara Benjamin for providing human PBMC samples including GVHD patients for flow cytometric analysis.

The authors wish to recognize the help and support of Paolo Serafini, PhD, Professor, Department of Microbiology and Immunology, and his lab for help with performing CODEX analysis using the Akoya.

The authors sincere appreciation is extended to Sarah Garce Lime for her invaluable technical support throughout this study, and Casey O. Lightbourn, Gina Adams, Jaret Fensterstock, and Alex Tudor for their work in the laboratory.

The researchers are indebted to Drs. Hyun Park (NCI, NIH) who kindly provided the TNFRSF25(DR3)-KO mice and Nurcin Liman for her coordination and effort in the experiments.

The authors would also like to thank Dr. Sophie Paczesny (Department of Microbiology and Immunology, Medical University of South Carolina, Charleston, South Carolina, USA.) for graciously providing the BALB/c MLL-AF9^GFP+^ cells used in the GVL studies and Heat Biologics, Inc for the mPTX-35 antibody.

## Authorship

Contribution: D.M., S.C., B.P., D.W., H.B., S.M., A.K., S.K., M.B.D.S, H.H., and M.G. performed research; M.R.M.v.d.B., M.T.A., G.R.H., and V.L.P provided reagents and support; D.M., S.C., B.P., H.B., M.B.D.S., H.H., and R.B.L. designed research and analyzed data; and D.M., S.C., B.P., D.W., and R.B.L wrote the manuscript. H.B., M.B.D.S, H.H, G.R.H. and V.L.P. critiqued the manuscript.

### Current Affiliations

Sabrina Copsel, PhD: Makana Therapeutics Miami, FL Seitaro Komai, MD, PhD: Kwantlen Polytechnic University, Kyoto, Japan.

Marcel RM van den Brink M.D., PhD: City of Hope, 1500 East Duarte Road, Duarte, CA 91010

Maria T. Abreu, MD: Cedars Sinai 110 George Burns Road, Davis Building D4063, Los Angeles, CA 90048. Phone: 310-423-7723.

## References

1. Bertaina A, Andreani M. Major Histocompatibility Complex and Hematopoietic Stem Cell Transplantation: Beyond the Classical HLA Polymorphism. Int J Mol Sci. 2018;19(2):621.

2. Choi SW, Levine JE, Ferrara JLM. Pathogenesis and Management of Graft-versus-Host Disease. Immunol Allergy Clin North Am. 2010;30(1):75–101.

3. Zeiser R. Advances in understanding the pathogenesis of graft-versus-host disease. Br J Haematol. 2019;187(5):563–572.

4. Hill GR, W. Krenger, J. L. Ferrara. The role of cytokines in acute graft-versus-host disease. Cytokines Cell Mol Ther. 1997;3(4):257–269.

5. Kumar S, Leigh ND, Cao X. The Role of Co-stimulatory/Co-inhibitory Signals in Graft-vs.-Host Disease. Front Immunol. 2018;9:.

6. Hamilton BK. Current approaches to prevent and treat GVHD after allogeneic stem cell transplantation. Hematology. 2018;2018(1):228–235.

7. Dou L, Zhao Y, Yang J, et al. Ruxolitinib plus steroids for acute graft versus host disease: a multicenter, randomized, phase 3 trial. Signal Transduct Target Ther. 2024;9(1):288.

8. Zhang M, Zhao P, Zhang Y, Wang J. Efficacy and safety of ruxolitinib for steroid-refractory graft-versus-host disease: Systematic review and meta-analysis of randomised and non-randomised studies. PLoS One. 2022;17(7):e0271979.

9. Zeiser R, von Bubnoff N, Butler J, et al. Ruxolitinib for Glucocorticoid-Refractory Acute Graft-versus-Host Disease. New England Journal of Medicine. 2020;382(19):1800–1810.

10. Raju G, Walji M, Nemirovsky D, et al. Real-World Experience Study of Belumosudil in Steroid-Refractory Chronic Graft-Versus-Host Disease (cGVHD) Demonstrated High Treatment Response without Significant Toxicities. Transplant Cell Ther. 2024;30(2):S277–S278.

11. Cutler C, Lee SJ, Arai S, et al. Belumosudil for chronic graft-versus-host disease after 2 or more prior lines of therapy: the ROCKstar Study. Blood. 2021;138(22):2278–2289.

12. Kean LS, Burns LJ, Kou TD, et al. Abatacept for acute graft-versus-host disease prophylaxis after unrelated donor hematopoietic cell transplantation. Blood. 2024;144(17):1834–1845.

13. Ngwube A, Rangarajan H, Shah N. Role of abatacept in the prevention of graft-versus-host disease: current perspectives. Ther Adv Hematol. 2023;14:20406207231152644.

14. Koshy AG, Kim HT, Liegel J, et al. Phase II clinical trial evaluating Abatacept in patients with steroid-refractory chronic graft versus host disease. Blood. 2023;

15. Wolf D, Barreras H, Bader CS, et al. Marked in Vivo Donor Regulatory T Cell Expansion via Interleukin-2 and TL1A-Ig Stimulation Ameliorates Graft-versus-Host Disease but Preserves Graft-versus-Leukemia in Recipients after Hematopoietic Stem Cell Transplantation. Biology of Blood and Marrow Transplantation. 2017;23(5):757–766.

16. Matsuoka K, Koreth J, Kim HT, et al. Low-Dose Interleukin-2 Therapy Restores Regulatory T Cell Homeostasis in Patients with Chronic Graft-Versus-Host Disease. Sci Transl Med. 2013;5(179):.

17. Ganguly S, Ross DB, Panoskaltsis-Mortari A, et al. Donor CD4+ Foxp3+ regulatory T cells are necessary for posttransplantation cyclophosphamide-mediated protection against GVHD in mice. Blood. 2014;124(13):2131–2141.

18. Hoffmann P, Ermann J, Edinger M, Fathman CG, Strober S. Donor-type CD4+CD25+ Regulatory T Cells Suppress Lethal Acute Graft-Versus-Host Disease after Allogeneic Bone Marrow Transplantation. Journal of Experimental Medicine. 2002;196(3):389–399.

19. Edinger M, Hoffmann P, Ermann J, et al. CD4+CD25+ regulatory T cells preserve graft-versus-tumor activity while inhibiting graft-versus-host disease after bone marrow transplantation. Nat Med. 2003;9(9):1144–1150.

20. Baron KJ, Turnquist HR. Clinical Manufacturing of Regulatory T Cell Products For Adoptive Cell Therapy and Strategies to Improve Therapeutic Efficacy. Organogenesis. 2023;19(1):.

21. Dong S, Hiam-Galvez KJ, Mowery CT, et al. The effect of low-dose IL-2 and Treg adoptive cell therapy in patients with type 1 diabetes. JCI Insight. 2021;6(18):.

22. Hippen KL, Hefazi M, Larson JH, Blazar BR. Emerging translational strategies and challenges for enhancing regulatory T cell therapy for graft-versus-host disease. Front Immunol. 2022;13:.

23. Bluestone JA, McKenzie BS, Beilke J, Ramsdell F. Opportunities for Treg cell therapy for the treatment of human disease. Front Immunol. 2023;14:.

24. Brunstein CG, Miller JS, Cao Q, et al. Infusion of ex vivo expanded T regulatory cells in adults transplanted with umbilical cord blood: safety profile and detection kinetics. Blood. 2011;117(3):1061–1070.

25. Matta BM, Reichenbach DK, Zhang X, et al. Peri-alloHCT IL-33 administration expands recipient T-regulatory cells that protect mice against acute GVHD. Blood. 2016;128(3):427–439.

26. Landwehr-Kenzel S, Müller-Jensen L, Kuehl J-S, et al. Adoptive transfer of ex vivo expanded regulatory T cells improves immune cell engraftment and therapy-refractory chronic GvHD. Molecular Therapy. 2022;30(6):2298–2314.

27. Sakaguchi S, Sakaguchi N, Shimizu J, et al. Immunologic tolerance maintained by CD25 ^+^ CD4 ^+^ regulatory T cells: their common role in controlling autoimmunity, tumor immunity, and transplantation tolerance. Immunol Rev. 2001;182(1):18–32.

28. Hanash AM, Levy RB. Donor CD4+CD25+ T cells promote engraftment and tolerance following MHC-mismatched hematopoietic cell transplantation. Blood. 2005;105(4):1828–1836.

29. Elias S, Rudensky AY. Therapeutic use of regulatory T cells for graft-versus-host disease. Br J Haematol. 2019;187(1):25–38.

30. Beauford SS, Kumari A, Garnett-Benson C. Ionizing radiation modulates the phenotype and function of human CD4+ induced regulatory T cells. BMC Immunol. 2020;21(1):18.

31. Qu Y, Jin S, Zhang A, et al. Gamma-Ray Resistance of Regulatory CD4 ^+^ CD25 ^+^ Foxp3 ^+^ T Cells in Mice. Radiat Res. 2010;173(2):148–157.

32. Anderson BE, McNiff JM, Matte C, et al. Recipient CD4+ T cells that survive irradiation regulate chronic graft-versus-host disease. Blood. 2004;104(5):1565–1573.

33. Balogh A, Persa E, Bogdándi EN, et al. The effect of ionizing radiation on the homeostasis and functional integrity of murine splenic regulatory T cells. Inflammation Research. 2013;62(2):201–212.

34. Ganguly S, Ross DB, Panoskaltsis-Mortari A, et al. Donor CD4+ Foxp3+ regulatory T cells are necessary for posttransplantation cyclophosphamide-mediated protection against GVHD in mice. Blood. 2014;124(13):2131–2141.

35. Fletcher RE, Nunes NS, Patterson MT, et al. Posttransplantation cyclophosphamide expands functional myeloid-derived suppressor cells and indirectly influences Tregs. Blood Adv. 2023;7(7):1117–1129.

36. Kanakry CG, Ganguly S, Zahurak M, et al. Aldehyde Dehydrogenase Expression Drives Human Regulatory T Cell Resistance to Posttransplantation Cyclophosphamide. Sci Transl Med. 2013;5(211):.

37. Strobl J, Pandey RV, Krausgruber T, et al. Long-term skin-resident memory T cells proliferate in situ and are involved in human graft-versus-host disease. Sci Transl Med. 2020;12(570):.

38. Divito SJ, Aasebø AT, Matos TR, et al. Peripheral host T cells survive hematopoietic stem cell transplantation and promote graft-versus-host disease. Journal of Clinical Investigation. 2020;130(9):4624–4636.

39. Bayer AL, Jones M, Chirinos J, et al. Host CD4+CD25+ T cells can expand and comprise a major component of the Treg compartment after experimental HCT. Blood. 2009;113(3):733–743.

40. Wang ECY, Thern A, Denzel A, et al. DR3 Regulates Negative Selection during Thymocyte Development. Mol Cell Biol. 2001;21(10):3451–3461.

41. Copsel SN, Garrido VT, Barreras H, et al. Minnelide suppresses GVHD and enhances survival while maintaining GVT responses. JCI Insight. 2024;9(9):.

42. Freeman D, Diefenbach C, Lam L, et al. terraFlow, a high-parameter analysis tool, reveals T cell exhaustion and dysfunctional cytokine production in classical Hodgkin’s lymphoma. Cell Rep. 2024;43(8):114605.

43. Khan SQ, Tsai MS, Schreiber TH, et al. Cloning, Expression, and Functional Characterization of TL1A-Ig. The Journal of Immunology. 2013;190(4):1540–1550.

44. Stein-Thoeringer CK, Nichols KB, Lazrak A, et al. Lactose drives *Enterococcus* expansion to promote graft-versus-host disease. Science (1979). 2019;366(6469):1143–1149.

45. Docampo MD, da Silva MB, Lazrak A, et al. Alloreactive T cells deficient of the short-chain fatty acid receptor GPR109A induce less graft-versus-host disease. Blood. 2022;139(15):2392–2405.

46. Burgos da Silva M, Ponce DM, Dai A, et al. Preservation of the fecal microbiome is associated with reduced severity of graft-versus-host disease. Blood. 2022;140(22):2385–2397.

47. Magnúsdóttir S, Heinken A, Kutt L, et al. Generation of genome-scale metabolic reconstructions for 773 members of the human gut microbiota. Nat Biotechnol. 2017;35(1):81–89.

48. Haak BW, Littmann ER, Chaubard J-L, et al. Impact of gut colonization with butyrate producing microbiota on respiratory viral infection following allo-HCT. Blood. 2018;blood-2018-01-828996.

49. Copsel S, Wolf D, Kale B, et al. Very Low Numbers of CD4+ FoxP3+ Tregs Expanded in Donors via TL1A-Ig and Low-Dose IL-2 Exhibit a Distinct Activation/Functional Profile and Suppress GVHD in a Preclinical Model. Biology of Blood and Marrow Transplantation. 2018;24(9):1788–1794.

50. Bayer AL, Chirinos J, Cabello C, et al. Expansion of a restricted residual host T _reg_ -cell repertoire is dependent on IL-2 following experimental autologous hematopoietic stem transplantation. Eur J Immunol. 2011;41(12):3467–3478.

51. Kanakry CG, Ganguly S, Zahurak M, et al. Aldehyde dehydrogenase expression drives human regulatory T cell resistance to posttransplantation cyclophosphamide. Sci Transl Med. 2013;5(211):211ra157.

52. Häcker G. GVHD prediction based on the microbiome. Blood. 2022;140(22):2313–2314.

53. Mathewson ND, Jenq R, Mathew A V, et al. Gut microbiome–derived metabolites modulate intestinal epithelial cell damage and mitigate graft-versus-host disease. Nat Immunol. 2016;17(5):505–513.

54. Masetti R, Zama D, Leardini D, et al. Microbiome-Derived Metabolites in Allogeneic Hematopoietic Stem Cell Transplantation. Int J Mol Sci. 2021;22(3):1197.

55. Copsel SN, Wolf D, Pfeiffer B, et al. Recipient Tregs: Can They Be Exploited for Successful Hematopoietic Stem Cell Transplant Outcomes? Front Immunol. 2022;13:.

56. Koreth J, Matsuoka K, Kim HT, et al. Interleukin-2 and Regulatory T Cells in Graft-versus-Host Disease. New England Journal of Medicine. 2011;365(22):2055–2066.

57. Bettini M, Bettini ML. Function, Failure, and the Future Potential of Tregs in Type 1 Diabetes. Diabetes. 2021;70(6):1211–1219.

58. Kukreja A, Cost G, Marker J, et al. Multiple immuno-regulatory defects in type-1 diabetes. Journal of Clinical Investigation. 2002;109(1):131–140.

59. Copsel SN, Malek TR, Levy RB. Medical Treatment Can Unintentionally Alter the Regulatory T-Cell Compartment in Patients with Widespread Pathophysiologic Conditions. Am J Pathol. 2020;190(10):2000–2012.

60. Honing DY, Luiten RM, Matos TR. Regulatory T Cell Dysfunction in Autoimmune Diseases. Int J Mol Sci. 2024;25(13):.

61. Wolf D, Schreiber TH, Tryphonopoulos P, et al. Tregs Expanded In Vivo by TNFRSF25 Agonists Promote Cardiac Allograft Survival. Transplantation. 2012;94(6):569–574.

62. Harris F, Berdugo YA, Tree T. IL-2-based approaches to Treg enhancement. Clin Exp Immunol. 2023;211(2):149–163.

63. Efe O, Gassen RB, Morena L, et al. A humanized IL-2 mutein expands Tregs and prolongs transplant survival in preclinical models. Journal of Clinical Investigation. 2024;134(5):.

64. Khoryati L, Pham MN, Sherve M, et al. An IL-2 mutein engineered to promote expansion of regulatory T cells arrests ongoing autoimmunity in mice. Sci Immunol. 2020;5(50):.

65. Schreiber TH, Wolf D, Tsai MS, et al. Therapeutic Treg expansion in mice by TNFRSF25 prevents allergic lung inflammation. Journal of Clinical Investigation. 2010;120(10):3629–3640.

66. McDonald-Hyman C, Flynn R, Panoskaltsis-Mortari A, et al. Therapeutic regulatory T-cell adoptive transfer ameliorates established murine chronic GVHD in a CXCR5-dependent manner. Blood. 2016;128(7):1013–1017.

67. Hippen KL, Merkel SC, Schirm DK, et al. Massive ex vivo expansion of human natural regulatory T cells (T(regs)) with minimal loss of in vivo functional activity. Sci Transl Med. 2011;3(83):83ra41.

68. Hoffmann P, Eder R, Boeld TJ, et al. Only the CD45RA+ subpopulation of CD4+CD25high T cells gives rise to homogeneous regulatory T-cell lines upon in vitro expansion. Blood. 2006;108(13):4260–4267.

69. Hoffmann P, Eder R, Kunz-Schughart LA, Andreesen R, Edinger M. Large-scale in vitro expansion of polyclonal human CD4+CD25high regulatory T cells. Blood. 2004;104(3):895–903.

70. Di Ianni M, Falzetti F, Carotti A, et al. Tregs prevent GVHD and promote immune reconstitution in HLA-haploidentical transplantation. Blood. 2011;117(14):3921–3928.

71. Wang ECY, Kitson J, Thern A, et al. Genomic structure, expression, and chromosome mapping of the mouse homologue for the WSL-1 ( DR3, Apo3, TRAMP, LARD, TR3, TNFRSF12 ) gene. Immunogenetics. 2001;53(1):59–63.

72. Fang L, Adkins B, Deyev V, Podack ER. Essential role of TNF receptor superfamily 25 (TNFRSF25) in the development of allergic lung inflammation. J Exp Med. 2008;205(5):1037–1048.

73. Lu S-Y, Liu K-Y, Liu D-H, Xu L-P, Huang X-J. High frequencies of CD62L+ naive regulatory T cells in allografts are associated with a low risk of acute graft-*versus* -host disease following unmanipulated allogeneic haematopoietic stem cell transplantation. Clin Exp Immunol. 2011;165(2):264–277.

74. Magenau JM, Qin X, Tawara I, et al. Frequency of CD4+CD25hiFOXP3+ Regulatory T Cells Has Diagnostic and Prognostic Value as a Biomarker for Acute Graft-versus-Host-Disease. Biology of Blood and Marrow Transplantation. 2010;16(7):907–914.

75. Rieger K. Mucosal FOXP3+ regulatory T cells are numerically deficient in acute and chronic GvHD. Blood. 2006;107(4):1717–1723.

76. Nishikii H, Kim B-S, Yokoyama Y, et al. DR3 signaling modulates the function of Foxp3+ regulatory T cells and the severity of acute graft-versus-host disease. Blood. 2016;128(24):2846–2858.

77. Ichikawa T, Hirahara K, Kokubo K, et al. CD103hi Treg cells constrain lung fibrosis induced by CD103lo tissue-resident pathogenic CD4 T cells. Nat Immunol. 2019;20(11):1469–1480.

78. Ross D, Jones M, Komanduri K, Levy RB. Antigen and Lymphopenia-Driven Donor T Cells Are Differentially Diminished by Post-Transplantation Administration of Cyclophosphamide after Hematopoietic Cell Transplantation. Biology of Blood and Marrow Transplantation. 2013;19(10):1430– 1438.

79. Polak K, Marchal P, Taroni C, et al. CD4+ regulatory T cells lacking Helios and Eos. Biochem Biophys Res Commun. 2023;674:83–89.

80. Sebastian M, Lopez-Ocasio M, Metidji A, et al. Helios Controls a Limited Subset of Regulatory T Cell Functions. The Journal of Immunology. 2016;196(1):144–155.

81. Kim H-J, Barnitz RA, Kreslavsky T, et al. Stable inhibitory activity of regulatory T cells requires the transcription factor Helios. Science (1979). 2015;350(6258):334–339.

82. Getnet D, Grosso JF, Goldberg M V., et al. A role for the transcription factor Helios in human CD4+CD25+ regulatory T cells. Mol Immunol. 2010;47(7–8):1595–1600.

83. Yu W, Ji N, Gu C, et al. Coexpression of Helios in Foxp3+ Regulatory T Cells and Its Role in Human Disease. Dis Markers. 2021;2021:1–9.

84. Thornton AM, Lu J, Korty PE, et al. Helios ^+^ and Helios ^−^ Treg subpopulations are phenotypically and functionally distinct and express dissimilar TCR repertoires. Eur J Immunol. 2019;49(3):398–412.

85. Peled JU, Gomes ALC, Devlin SM, et al. Microbiota as Predictor of Mortality in Allogeneic Hematopoietic-Cell Transplantation. New England Journal of Medicine. 2020;382(9):822–834.

86. Jenq RR, Taur Y, Devlin SM, et al. Intestinal Blautia Is Associated with Reduced Death from Graft-versus-Host Disease. Biology of Blood and Marrow Transplantation. 2015;21(8):1373–1383.

87. Staffas A, Burgos da Silva M, van den Brink MRM. The intestinal microbiota in allogeneic hematopoietic cell transplant and graft-versus-host disease. Blood. 2017;129(8):927–933.

88. Furusawa Y, Obata Y, Fukuda S, et al. Commensal microbe-derived butyrate induces the differentiation of colonic regulatory T cells. Nature. 2013;504(7480):446–450.

89. He L, Zhong Z, Wen S, et al. Gut microbiota-derived butyrate restores impaired regulatory T cells in patients with AChR myasthenia gravis via mTOR-mediated autophagy. Cell Communication and Signaling. 2024;22(1):215.

90. Maslowski KM, Vieira AT, Ng A, et al. Regulation of inflammatory responses by gut microbiota and chemoattractant receptor GPR43. Nature. 2009;461(7268):1282–1286.

91. Burgueño JF, Fritsch J, Santander AM, et al. Intestinal Epithelial Cells Respond to Chronic Inflammation and Dysbiosis by Synthesizing H2O2. Front Physiol. 2019;10:.

92. Grasberger H, Gao J, Nagao-Kitamoto H, et al. Increased Expression of DUOX2 Is an Epithelial Response to Mucosal Dysbiosis Required for Immune Homeostasis in Mouse Intestine. Gastroenterology. 2015;149(7):1849–1859.

93. Sommer F, Bäckhed F. The gut microbiota engages different signaling pathways to induce Duox2 expression in the ileum and colon epithelium. Mucosal Immunol. 2015;8(2):372–379.

94. Castrillón-Betancur JC, López-Agudelo VA, Sommer N, et al. Epithelial Dual Oxidase 2 Shapes the Mucosal Microbiome and Contributes to Inflammatory Susceptibility. Antioxidants. 2023;12(10):1889.

95. Hazime H, Ducasa G, Fernandez I, et al. EPITHELIAL DUOX2 ACTIVATION INCREASES GUT PERMEABILITY AND BACTERIAL TRANSLOCATION THAT IS RESCUED WITH BUTYRATE SUPPLEMENTATION. Gastroenterology. 2022;162(3):S49.

96. Hazime H, Ducasa G, Brito N, et al. LOSS OF EPITHELIAL DUOX2 SIGNALING IS ACCOMPANIED BY A REDUCTION IN AKKERMANSIACEAE AND PROMOTES THE DEVELOPMENT OF METABOLIC SYNDROME IN A MICROBIOME-DEPENDENT MANNER. Inflamm Bowel Dis. 2023;29(Supplement_1):S72–S72.

97. Hill GR, Ferrara JLM. The primacy of the gastrointestinal tract as a target organ of acute graft-versus-host disease: rationale for the use of cytokine shields in allogeneic bone marrow transplantation. Blood. 2000;95(9):2754–2759.

