## Supplementary Materials for "Pretransplant targeting of TNFRSF25 and CD25 stimulates recipient Tregs in target tissues ameliorating GVHD post-HSCT"

### Supplemental Methods

#### Isolation of Colonic Epithelial Cells and Measurement of Epithelial H2O2 Production

Colonic and ileum sections were shaken at 80 rpm for 60 mins at room temperature in a chelation buffer prepared with 20 mM EDTA in HBSS. The residual EDTA is then gently washed with HBSS. Crypts were released by gentle agitation for 1 hour. To measure $H_2O_2$ rate, freshly isolated colonic epithelial cells were seeded in 96 well plates and incubated in Dulbecco's PBS (DPBS) solution containing Ca2+, Mg2+, 0.1 U/mL HRP (Sigma Aldrich), 30 μM 10-Acetyl-3,7-dihydroxyphenoxazine (Biotium), and 100 μM PMSF (Sigma-Aldrich). The real-time formation of fluorescent resorufin (Ex530 nm/Em 590 nm) was read at 60s intervals for 15 min at 37°C in a Synergy H1 fluorometer (BioTek). Immediately after $H_2O_2$ measurement, MTT (ATCC) assay was performed (per manufacturer's instructions) to normalize $H_2O_2$ production to the number of viable cells. The amount of $H_2O_2$ was determined using a standard curve of $H_2O_2$ prepared fresh for each experiment. All samples were assayed in triplicates.

#### Intestinal Barrier Analysis

Protocol was adapted from Hazime et. al. Transplanted mice underwent food restriction for 12 hours before receiving a 600-mg/kg oral gavage of 4 kilodaltons fluorescein isothiocyanate–dextran. Four hours later, blood was collected and fluorescein isothiocyanate fluorescence (485/528 nm ex/em) in separated plasma was read in a Synergy H1 fluorometer (BioTek). All samples were assayed in triplicates.[1]

#### Histopathology and Immunohistochemistry

Colon, ear skin, and liver were collected, formalin-fixed and paraffin-embedded. Sections were stained with Hematoxylin-Eosin (H&E) or Masson Trichrome for histologic examination. Slides were scored as previously described.[2] Sections were given a pathology score of 0–2 (0=normal, 1=moderate, 2=severe) based upon the amount of inflammation/infiltration, collagen deposition and dermal thickening for skin, and inflammation/infiltration, edema, mucosal thickening, and crypt structure for colon. Scores were then aggregated to calculate an overall histopathology score (max score = 6 for skin, 8 for colon, and liver).

**Isolating Lamina Propria lymphocytes from colons for phenotyping cell populations.**

Briefly modified from (Miltenyi Biotec Lamina Propria Dissociation kit). Intestines were removed and placed in HBSS (w/o) in a Petri dish. Feces were cleared by flushing with HBSS(w/o) using a syringe. The intestines were first cut longitudinally and then laterally into pieces of approximately 0.5 cm length. Tissue pieces were transferred into a 50 ml tube containing 20 ml of pre-digestion solution: (1xHBSS(w/o) containing 5 mM EDTA, 5% fetal bovine serum (FBS), 1 mM DTT). Samples were incubated for 20 minutes at 37 C under continuous rotation. Samples were mixed well and applied to a 100 uM mesh filter placed on a 50 ml collection tube. Lamina propria tissue pieces were transferred into a new 50 ml tube containing 20 ml of fresh pre-digestion solution and the incubation repeated. Lamina propria pieces were then transferred into a new 50 ml tube containing 20 ml of HBSS (w/o) and incubated for 20 minutes at 37 C under continuous rotation. After a quick vortex, samples were applied to a 100-uM mesh filter placed on a 50 ml collection tube. A digestion solution containing 10% FBS Dnase1 @ 0.1 mg/ml and collagenase 3 @ 600 U/ml was prepared and preheated. Intestinal tissue was placed into a gentleMACS C tube containing 3 ml of digestion solution. Samples were incubated for 30 minutes at 37 C under continuous rotation. A C tube was attached upside down onto the sleeve of the gentleMACS Dissociator and the gentleMACS Program “m_intestine_01” run. After a short spin, samples were resuspended in 5 ml of phosphate buffer solution and applied to a 100 um mesh filter. The cell suspension was centrifuged at 300Xg for 10 minutes at room temperature, aspirated and lamina propria lymphocytes resuspended with appropriate buffer and volume for further applications.

**Isolating Lung lymphocytes for phenotyping cell populations**

Briefly modified from (Miltenyi Biotec Lung Dissociation kit). Lungs were perfused through the heart with 10 mL of 1X PBS. Lobes were harvested and placed on a petri dish with 1 mL PBS. Lungs were then transferred into GentleMACS C tubes with 3 mL of 1X PBS containing 2 mg/ml Collagenase Type IV (ThermoFisher, Gibco) and 1 mg/ml DNase (Sigma-Aldrich). The samples were placed on the GentleMACS Dissociator and run on cycle "m_lung_01". Afterwards, they were incubated at 37 C for 30 minutes with quick vortex every 5-minute intervals. Samples were then placed on the GentleMACS Dissociator on cycle "m_lung_02". Lung tissue was then poured over 100uM mesh cell strainer and into a collection tube. Samples were spun for 5 minutes at 1500 RPM and resuspended in 1 mL of ACK lysis buffer for 5 min at room temperature. The ACK buffer was then washed from the cells with 5 mL of 1X PBS. Cells were then resuspended with appropriate buffer and volume for further applications.

**Conjunctiva and lacrimal gland dissociation**

Conjunctival tissues from both eyes of each mouse were harvested at the study endpoint, carefully cut into small pieces, placed in 2.5mg/ml collagenase (Liberase, TL research grade, Roche) in 500μl of RPMI without serum and incubated at 37 ℃.[3] The tissues were pipetted to facilitate dissociation every 15 minutes, with the process repeated four times for a total incubation period of 60 minutes. A similar protocol was followed for the lacrimal glands, with the only difference being an incubation time of 45 minutes with Liberase. After digestion, the samples were passed through 40um cell strainers (BD Falcon, Franklin Lakes, NJ, USA) to obtain a single-cell suspension. The collagenase was inactivated using 1.5 ml of 10% fetal bovine serum (FBS) in RPMI. Cell counts and viability were assessed using a hemocytometer after trypan blue staining, followed by staining with the designated flow panel.

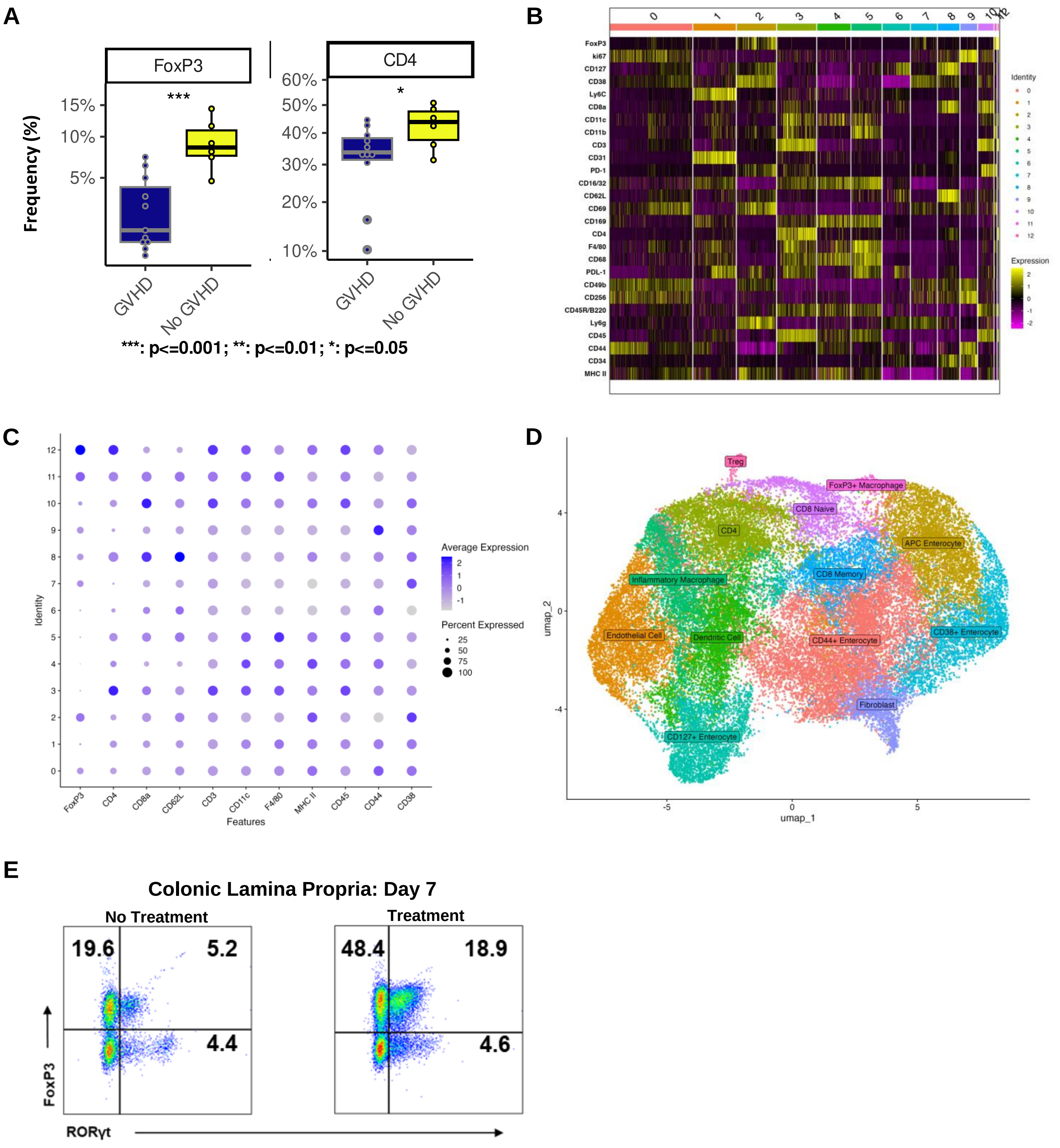


Supplemental Figure 1

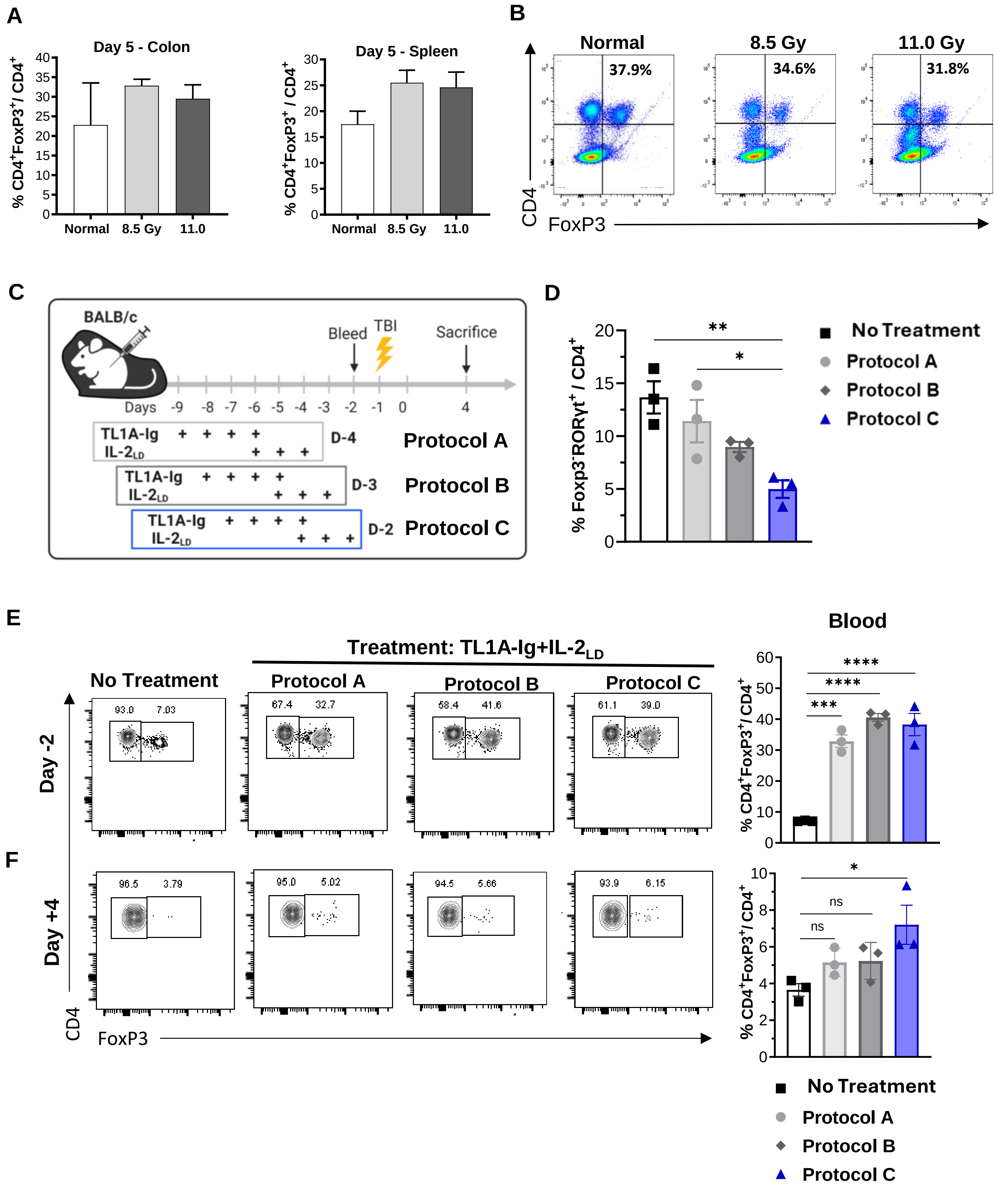


Supplemental Figure 2

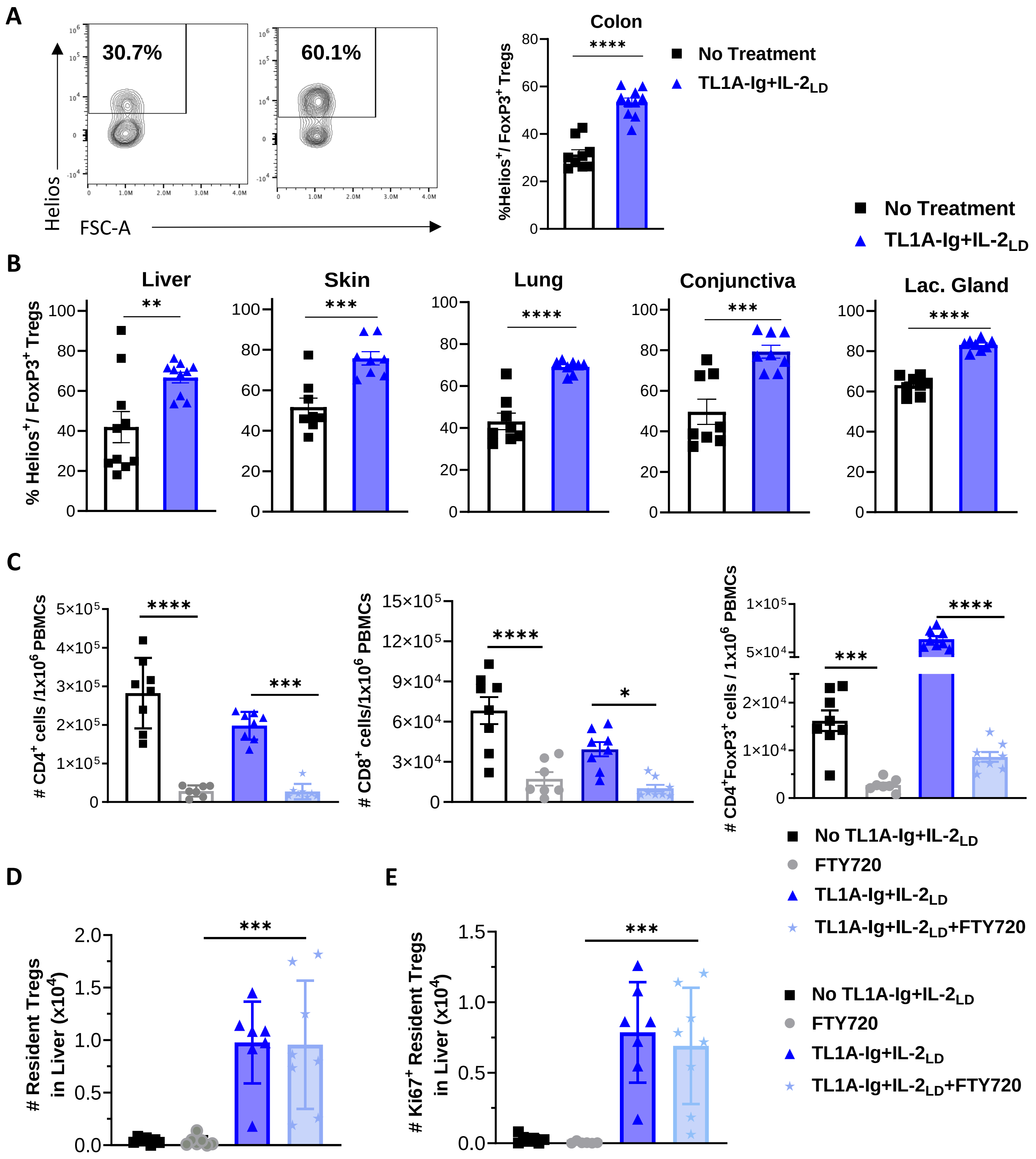


Supplemental Figure 3

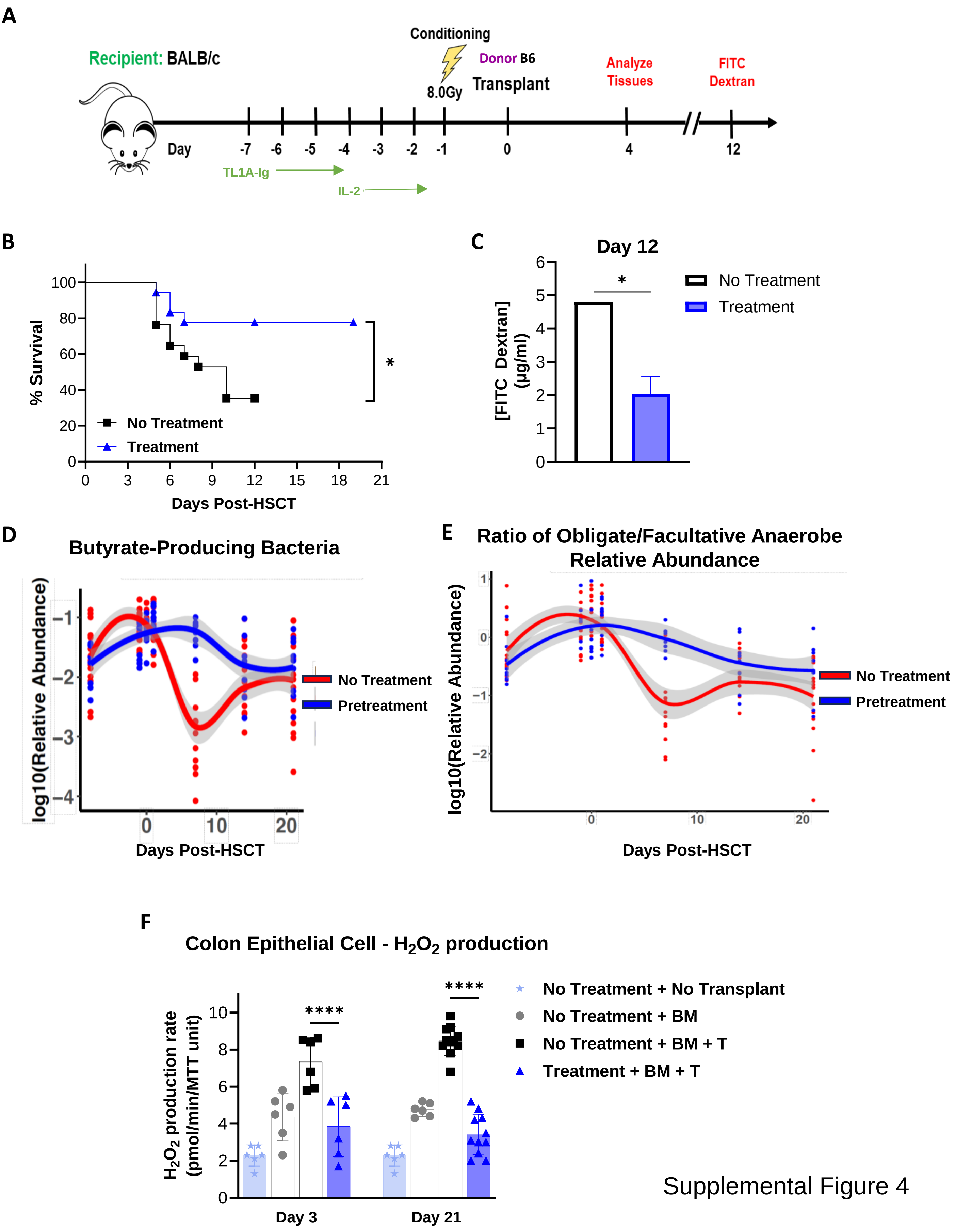


Supplemental Figure 4

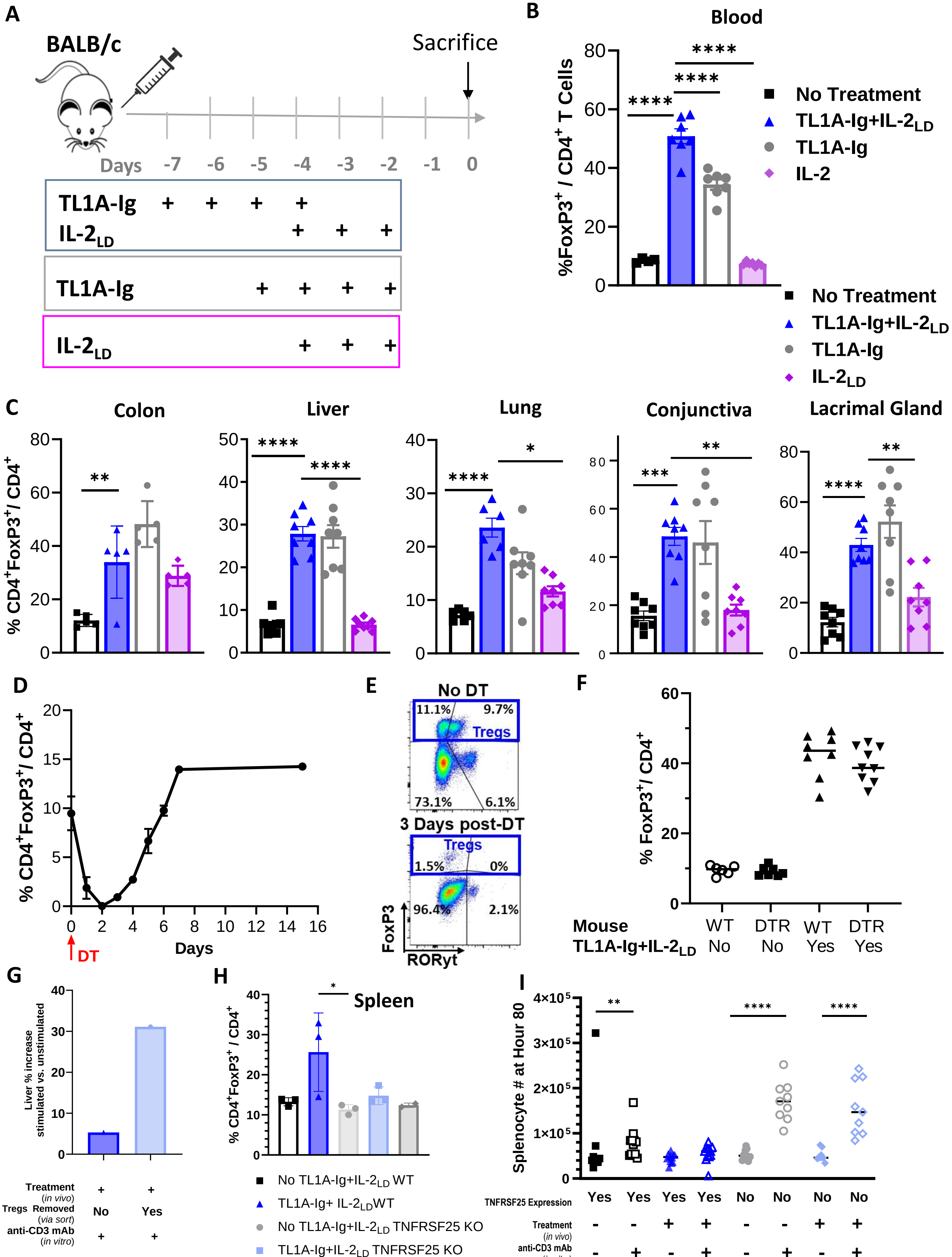


Supplemental Figure 5

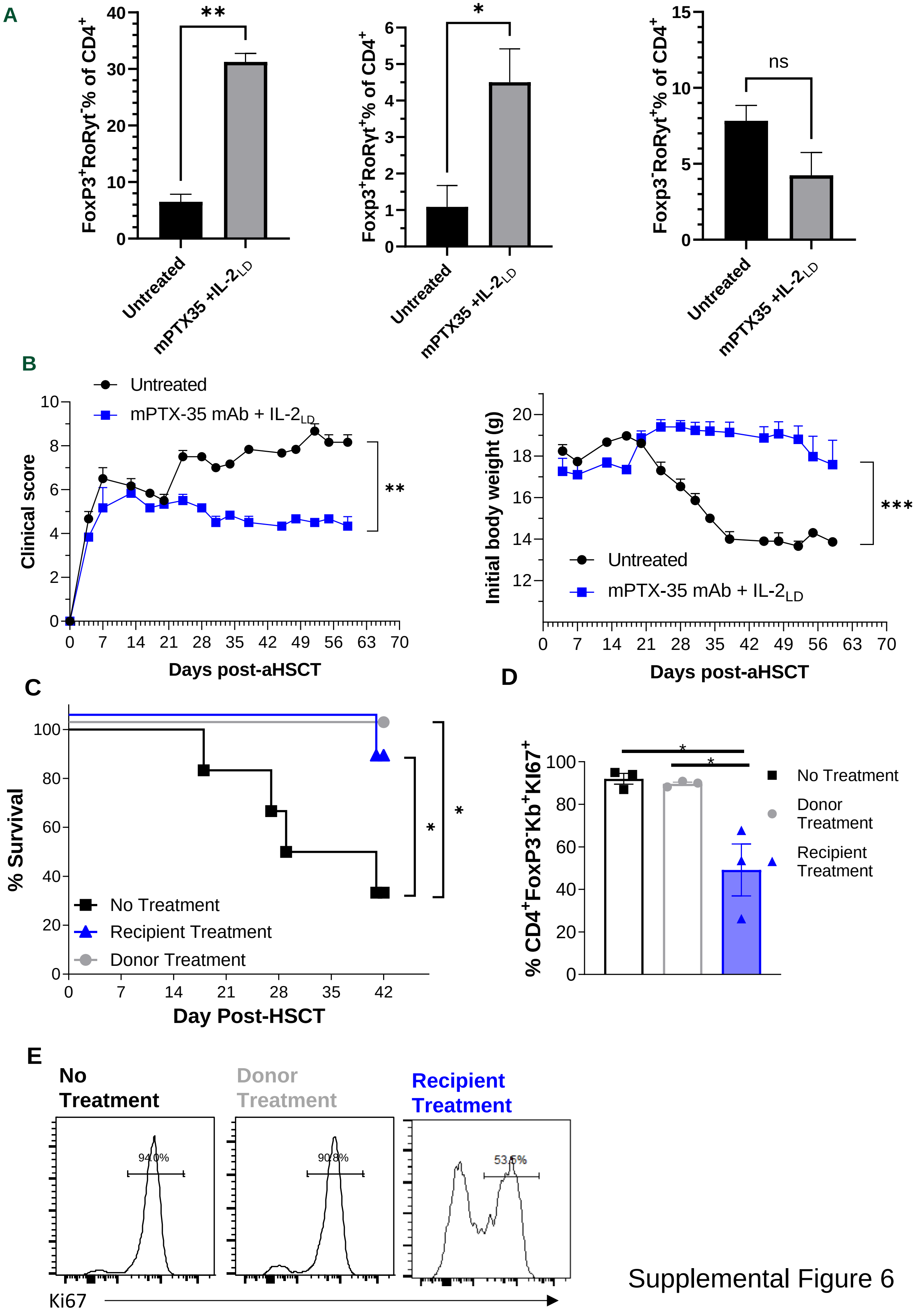


Supplemental Figure 6

**Supplemental Figure 1**

**Analysis of Tregs in patients and experimental mice with GVHD. (A)** Frequencies (%) of $CD4^+/FoxP3^+$ and $CD4^+$ T cells from human HSCTx recipients greater than 1 year post transplantation stratified by No GVHD (n=6, blue boxplots) and GVHD (n=11, yellow boxplots) determined by terraFlow (see methods). **(B)** Heat map of all 45,495 segmented cells showing the clusters based upon 27 markers. The width of the upper x-axis clusters represents the proportion of cells within each cluster. The individual black, purple or yellow lines represent all 27 markers expressed for each individual cell. Clustering was determined by PCA. **(C)** The dot plot represents the average expression of selected characteristic immune markers (gray to purple gradient) and the size of the dot represents the percent of cells expressing the marker within each cluster. This determined the type of cells within each cluster. **(D)** UMAP graphic showing all 45,495 cells after identifying and renaming each cluster group. Multiple subsets of enterocytes and macrophages were identified in the colon of BALB/c recipients at Day+24. This included a non-hematopoietic enterocyte population positive for MHCII. Data represents whole colonic tissue, n=2/group. **(E)** BALB/c mice were treated with TL1A-Ig+$IL2_{LD}$. Expansion of BALB/c Tregs from colon lamina propria at Day 7 (see Methods, TL1A-Ig+$IL2_{LD}$ *in vivo* administration), representative dot plots from individual mice are shown (n=5/group – see Fig. 1D).

**Supplemental Figure 2**

**Persistence of Tregs post-conditioning and development of TL1A-Ig+IL2$_{LD}$ treatment protocol pre-aHSCT**. **(A-B)** To demonstrate Treg persistence post-TBI conditioning, B6 FIR mice were either unmanipulated, i.e. normal (n=4/group) or irradiated with dosages of 8.5 or 11.0 Gy (n=2/group). Five days later, mice were sacrificed and Treg levels were analyzed from colon lamina propria and spleen. **(A)** Frequency of Tregs in colon and spleen at Day+5 post radiation. **(B)** Representative flow plots of Tregs in colon lamina propria at Day+5 post radiation. **(C-F)** BALB/c mice were treated as indicated with TL1A-Ig+IL2$_{LD}$ as illustrated in panel **(C)** at different time points and irradiated (TBI:8.5 Gy) on Day-1**. (D)** All mice were bled at Day-2 and the frequency of TH17 T cells were analyzed. **(E&F)** Treg frequencies at Day-2 **(E)** and Day+4 **(F)** are shown providing dot plots and summarized bar graphs.

**Supplemental Figure 3**

**Detection of Helios$^{+}$ Tregs after TL1A-Ig+IL2$_{LD}$ treatment in multiple tissues targeted during GVHD and identification of proliferating CD103$^{+}$CD69$^{+}$ tissue resident hepatic Tregs**. BALB/c mice were either left untreated or were treated with TL1A-Ig+IL2$_{LD}$. On Day8 colon, liver, skin, lung and ocular adnexa (conjunctiva and lacrimal gland) were harvested, homogenized and single cell suspensions made for analysis via flow cytometry. **(A)** Helios expression in Tregs of untreated and TL1A-Ig+IL2$_{LD}$ treated mice showing contour plots and summary bar graph of from colon lamina propria lymphocytes at Day8. **(B)** Frequency of Helios expression in FoxP3$^{+}$ Tregs was significantly higher in non-hematopoietic tissue compartments at Day8 of TL1A-Ig+IL2$_{LD}$ treated mice. **(C)** The S1PR modulator, FTY720, was used to delineate whether TL1A-Ig+IL2$_{LD}$ expanded Tregs are trafficking from hematopoietic sites or proliferating in non-hematopoietic tissues. PBMCs demonstrate a significant decrease in CD4$^{+}$ total, CD8$^{+}$, and CD4$^{+}$FoxP3$^{+}$ cell numbers per 1x10$^{6}$ PMBCs associated with FTY720 administration alone or with TL1A-Ig+IL2$_{LD}$ expansion. **(D)** Number of resident Tregs (CD69$^{+}$CD103$^{+}$) from liver show TL1A-Ig+IL2$_{LD}$ expansion independent of FTY720 indicating resident Treg proliferation. **(E)** Absolute numbers of proliferating (Ki67$^{+}$) resident Tregs (CD69$^{+}$CD103$^{+}$) in liver confirming marked Ki67 expression associated with TL1A-Ig+IL2$_{LD}$ expansion (N=8; compiled from >2 independent experiments). Data represent the mean ±SEM. *p<0.05; **p<0.01; ****p<0.0001

Supplemental Figure 4

**Pre-conditioning of recipients targeting TNFRSF25 and CD25 with TL1A-Ig+IL-$2_{LD}$ augments early survival following MHC-mismatched aHSCT and improves GI integrity post-transplant. (A)** BALB/c mice were pre-treated with TL1A-Ig+IL$2_{LD}$ as previously described prior to transplant with B6 donor. CD45.1 B6 ($H2^b$) TCD-BM ($5.5x10^6$) and $6.0x10^5$ splenic T cells were transplanted into BALB/c ($H2^d$) recipients on Day0 after TBI (8.0 Gy) on Day-1. **(B)** Survival curve showing increased survival with TL1A-Ig+IL$2_{LD}$ pretreatment. **(C)** FITC-dextran serum assessment on Day+12 showing less gastrointestinal permeability. Data represents a pool of two experiments, (n=2/experiment). FITC-dextran assessment of untreated mice represents the level obtained after pooling serum from this group (n=4). **(D,E)** Microbiome assessment from BALB/c mice after B6→BALB/c aHSCT (TBI:7.25 Gy) with one group receiving TL1A-Ig+IL-$2_{LD}$ pretreatment. Data presented was pooled from 2 experiments n=16 mice/group. **(D)** Relative abundance of Butyrate-producing bacteria and **(E)** Ratio of obligate/facultative anaerobe relative abundance from TL1A-Ig+IL-$2_{LD}$ pretreated and untreated mice over time (see Methods for detailed days). **(F)** To assess Duox2 function, we compared $H_2O_2$ production rates of colonic epithelial cells isolated from no transplant, BM only (BM), untreated (BM+T), and (BM+T) TL1A-Ig+IL-$2_{LD}$ pretreated mice on Days+3 and +21.

**Supplemental Figure 5**

**In GVHD targeted tissues, individual TL1A-Ig - but not IL-$2_{LD}$ - administration induces expansion of Tregs comparable to TL1A-Ig+IL-$2_{LD}$ and requires TNFRSF25 expression. (A)** Schematic administration of TL1A-Ig and/or IL-2LD to BALB/c mice assessing $CD4^{+}FoxP3^{+}$ cell in blood and non-hematopoietic tissues. Mice were sacrificed and tissues harvested on Day0. **(B)** On Day-2, animals were bled and frequency of $CD4^{+}FoxP3^{+}$ cells is shown. **(C)** $CD4^{+}FoxP3^{+}$ cell frequency in colon, liver, lung, and ocular adnexa (conjunctiva and lacrimal gland) at Day0. **(D-F)** BALB/$c^{FoxP3\text{-}DTR\text{–}eGFP}$ (DTR) mice were injected with 1 µg Diphtheria Toxin (DT) and Tregs levels were evaluated. **(D)** Treg peripheral blood levels were transiently decreased after injecting DT and by Day6 Treg levels returned to normal levels. **(E)** Representative dot plots showing peripheral blood Treg $FoxP3^{+}$ levels before DT and 3 days after DT injections. **(F)** TL1A-Ig+IL-$2_{LD}$ induces comparable Treg expansion in BALB/$c^{FoxP3\text{-}DTR\text{–}eGFP}$ mice and wild type BALB/c mice (unexpanded BALB/c, n=6; unexpanded DTR, n=7; expanded BALB/c, n=8; expanded DTR, n=9). **(G)** B6-$FoxP3^{RFP}$ mice were treated with TL1A-Ig+IL-$2_{LD}$ and liver $RFP^{+}$ Tregs were sorted. All $RFP^{-}$ cells from the sorting were plated with and without 1µg of anti-CD3 mAb, cultured for 120 hours, and then counted to evaluate proliferation comparing counts to the unstimulated (no anti-CD3) cultures for baseline proliferation levels. Each bar represents one experiment with 3 replicated wells that were pooled and counted. **(H)** TNFRSF25 expression is required for TL1A-Ig induced Treg expansion *in vivo*. Frequency of $CD4^{+}FoxP3^{+}$ Tregs in spleens at Day0 from their respective mice (BALB/c and BALB/c-TNFRSF25KO) ± TL1A-Ig+IL-$2_{LD}$. **(I)** Loss of suppression in the absence of the expression of TNFRSF25 followed by TL1A-Ig in vivo stimulation. Number of total splenocytes at 80 hours after culture by their respective mice, TL1A-Ig+IL-$2_{LD}$ treatment, and anti-CD3 mAb (1µg). Data represents the mean ±SEM*$p<0.05$; **$p<0.01$; ***$p<0.001$ ****$p<0.0001$.

**Supplemental Figure 6**

**Donor and recipient expanded Tregs significantly diminished GVHD severity. (A-B)** BALB/c mice were treated before TBI with an anti-TNFRSF25 agonistic mAb and IL-2 (mPTX-35+IL-$2_{LD}$). Allo-HSCT was performed (B6→BALB/c) with 5 mice/group. **(A)** Four days post-aHSCT colonic lamina propria Tregs were elevated in treated mice versus untreated with no difference in TH17 cell frequency. **(B)** GVHD clinical score and body weight percentage post-aHSCT. **(C-E)** B6 donor or recipient BALB/c mice were TL1A-Ig+IL-$2_{LD}$ treated before TBI (7.5 Gy). There were 8 mice/group. **(C)** Donor or recipient treatment show increased overall survival. **(D)** Summary bar graph showing reduced splenic donor $CD4^+FoxP3^-Ki67^+$ (T conv) proliferation frequency in TL1A-Ig+IL-$2_{LD}$ pretreated recipients versus untreated and donor TL1A-Ig+IL-$2_{LD}$ treatment on Day +5. **(E)** Representative histograms illustrating splenic donor T conv cells Ki67 expression showing Day 5 post-aHSCT donor Tconv $Ki67^+$ splenic cells. Data represent the mean ±SEM*p<0.05; **p<0.01; ***p<0.001 ****p<0.0001.

### Conventional Flowcytometery Antibody Materials Table 1

| Species | Marker | Clone | Flurophore | Ab Source - Company | Catologue Number |
|---|---|---|---|---|---|
| Mouse | CD44 | IM7 | BUV 737 | BD Biosciences | 612799 |
| Mouse | Ly6c | HK1.4 | PE Dazzle 594 | Biolegend | 128044 |
| Mouse | Ly6c | AL-21 | FITC | BD Biosciences | 553104 |
| Mouse | Viability | Live/Dead Blue | N/A | Invitrogen | L23105 |
| Mouse | KLRG1 | 2F1 | BV 605 | Biolegend | 138419 |
| Mouse | FoxP3 | FJK-16s | Alexa Fluor 700 | Invitrogen | 56-5773-82 |
| Mouse | CD62L | MEL-14 | FITC | BD Biosciences | 553150 |
| Mouse | ICOS | 7E.17G9 | PE-Cy5 | Invitrogen | 15-9949-82 |
| Mouse | CD25 | PC61 | PE | Biolegend | 102008 |
| Mouse | RORgt | Q31-378 | Alexa Fluor 647 | BD Biosciences | 562682 |
| Mouse | CD4 | GK1.5 | BUV 805 | BD Biosciences | 612900 |
| Mouse | CD8 | 53-6.7 | BUV 661 | BD Biosciences | 569186 |
| Mouse | Helios | 22F6 | eF 450 | Invitrogen | 48-9883-42 |
| Mouse | CD45.2 | 104 | BUV 395 | Invitrogen | 363-0454-82 |
| Mouse | CD3 | 17A2 | RB780 | BD Biosciences | 755792 |
| Mouse | DR3 | 4C12 | PE | Biolegend | 144406 |
| Mouse | Ki-67 | SolA15 | PECy7 | Invitrogen | 25-5698-82 |
| Mouse | CD103 | 2E7 | BUV 615 | BD Biosciences | 751631 |
| Mouse | CD69 | H1.2F3 | FITC | Biolegend | 104506 |
| Human | Live/Dead | N/A | UV Blue | Invitrogen | L23105A |
| Human | CD45RA | HI100 | BUV395 | BD Biosciences | 740298 |
| Human | CD69 | FN50 | BUV563 | BD Biosciences | 748764 |
| Human | CD4 | RPA-T4 | BUV615 | BD Biosciences | 751347 |
| Human | CD95 | DX2 | BUV737 | BD Biosciences | 612790 |
| Human | CD8 | SK1 | BUV805 | BD Biosciences | 612889 |
| Human | CD27 | O323 | BV421 | BioLegend | 302824 |
| Human | CD19 | SJ25C1 | Pacific Blue | BioLegend | 363036 |
| Human | Granzyme B | GB11 | BV510 | BD Biosciences | 563388 |
| Human | CD3e | UCHT1 | BV570 | BioLegend | 300436 |
| Human | CD28 | CD28.2 | BV605 | BioLegend | 302968 |
| Human | CD127 | A019D5 | BV711 | BioLegend | 351327 |
| Human | Ki67 | Ki67 | BV750 | BioLegend | 350536 |
| Human | CTLA-4 | BNI3 | BV785 | BioLegend | 369624 |
| Human | CX3CR1 | 2A9-1 | FITC | BioLegend | 341606 |
| Human | CD39 | A1 | PE Dazzle594 | BioLegend | 328224 |
| Human | CD45RO | UCHL1 | PE Cy5 | Invitrogen | 15-0457-42 |
| Human | CD25 | M-A251 | PE Fire700 | BioLegend | 356145 |
| Human | PD-1 | EH12.2H7 | PE Cy7 | BioLegend | 329918 |
| Human | CXCR3 | 6025H7 | PE Fire810 | BioLegend | 353760 |
| Human | TOX | REA473 | APC | MACS Miltenyi Biotec | 130-118-335 |
| Human | TCF-1 | S33-966 | AF647 | BD Biosciences | 266693 |
| Human | FoxP3 | PCH101 | AF700 | Invitrogen | 56-4776-41 |
| Human | CCR7 | G043H7 | APC Fire810 | BioLegend | 353264 |
| Mouse | CD | RM4-5 | PE | Biolegend | 100512 |

### Spatial Biology Materials Table 2

| Marker | BarCode | ReporterCode | Exposure Time (ms) | Ab Source - Company | Catologue Number |
|---|---|---|---|---|---|
| CD31 | BX002 | RX002 | 200 | Akoya | 4250001 |
| CD44 | BX005 | RX005 | 150 | Akoya | 4250002 |
| CD45 | BX007 | RX007 | NA | Akoya | 4150002 |
| CD45R/B220 | BX010 | RX010 | 100 | Akoya | 4150006 |
| FoxP3 | BX013 | RX013 | 200 | eBioscience | 14-4777-82 |
| MHCII | BX014 | RX014 | 100 | Akoya | 4250003 |
| CD169 | BX015 | RX015 | 100 | Akoya | 4550100 |
| CD69 | BX016 | RX016 | 150 | BioLegend | 104502 |
| CD62L | BX017 | RX017 | 200 | BioLegend | 104402 |
| CD38 | BX019 | RX019 | 100 | Akoya | 4150013 |
| CD3 | BX021 | RX021 | 150 | Akoya | 4550109 |
| CD127 | BX023 | RX023 | 100 | BioLegend | 135002 |
| Ly6G | BX024 | RX024 | 90 | Akoya | 4550110 |
| CD11b | BX025 | RX025 | 200 | Akoya | 4150015 |
| CD4 | BX026 | RX026 | 200 | Akoya | 4250016 |
| Ly6C | BX027 | RX027 | 150 | BioLegend | 128001 |
| CD8a | BX029 | RX029 | 200 | Akoya | 4250017 |
| CD11c | BX030 | RX030 | 150 | Akoya | 4550108 |
| PD1 | BX031 | RX031 | 200 | BioLegend | 135202 |
| PDL1 | BX032 | RX032 | 150 | BioLegend | 124302 |
| CD256 | BX035 | RX035 | 200 | ThermoFisher | PA5-86370 |
| CD16/32 | BX036 | RX036 | 100 | BD | 553142 |
| F4/80 | BX037 | RX037 | 200 | BD | 565409 |
| CD49b | BX040 | RX040 | 150 | BioLegend | 108902 |
| CD68 | BX042 | RX042 | 90 | AbCam | ab53444 |
| CD25 | BX046 | RX046 | 200 | BioLegend | 154202 |
| Ki67 | BX047 | RX047 | 200 | Akoya | 4250019 |

Classification of Bacterial Organisms by Oxygen Metabolism Table 3

| Classification | Oxygen Metabolism Groups |
|---|---|
| Strict anaerobes | Anaerobes |
| Strict anaerobes | Strict anaerobes |
| Tolerant anaerobes | Facultative anaerobes |
| Tolerant anaerobes | Aerotolerants |
| Tolerant anaerobes | Microaerophile |
| Tolerant anaerobes | Obligate aerobes |
| Tolerant anaerobes | Aerobes |

**Supplemental Materials References**


1. Hazime, H. *et al.* Intestinal Epithelial Inactivity of Dual Oxidase 2 Results in Microbiome-Mediated Metabolic Syndrome. *Cell Mol Gastroenterol Hepatol* 16, 557–572 (2023).
2. Kaplan, D. H. *et al.* Target Antigens Determine Graft-versus-Host Disease Phenotype. *The Journal of Immunology* 173, 5467–5475 (2004).
3. Perez, V. L. *et al.* Meibomian Gland Dysfunction: A Route of Ocular Graft-Versus-Host Disease Progression That Drives a Vicious Cycle of Ocular Surface Inflammatory Damage. *Am J Ophthalmol* 247, 42–60 (2023).
